# Patterns and Mechanisms of Sex Ratio Distortion in the Collaborative Cross Mouse Mapping Population

**DOI:** 10.1101/2021.06.23.449644

**Authors:** Brett A. Haines, Francesca Barradale, Beth L. Dumont

## Abstract

In species with single-locus, chromosome-based mechanisms of sex determination, the laws of segregation predict an equal ratio of females to males at birth. Here, we show that departures from this Mendelian expectation are commonplace in the 8-way recombinant inbred Collaborative Cross (CC) mouse population. More than one-third of CC strains exhibit significant sex ratio distortion (SRD) at wean, with twice as many male-biased than female-biased strains. We show that these pervasive sex biases persist across multiple breeding environments, are stable over time, are not fully mediated by maternal effects, and are not explained by sex-biased neonatal mortality. SRD exhibits a heritable component, but QTL mapping analyses and targeted investigations of sex determination genes fail to nominate any large effect loci. These findings, combined with the reported absence of sex ratio biases in the CC founder strains, suggest that SRD manifests from multilocus combinations of alleles only uncovered in recombined CC genomes. We speculate that the genetic shuffling of eight diverse parental genomes during the early CC breeding generations led to the decoupling of sex-linked drivers from their co-evolved suppressors, unleashing complex, multiallelic systems of sex chromosome drive. Consistent with this interpretation, we show that several CC strains exhibit copy number imbalances at co-evolved X-and Y-linked ampliconic genes that have been previously implicated in germline genetic conflict and SRD in house mice. Overall, our findings reveal the pervasiveness of SRD in the CC population and nominate the CC as a powerful resource for investigating sex chromosome genetic conflict in action.

**ARTICLE SUMMARY:** We compiled breeding records from The Collaborative Cross (CC) mouse mapping population to quantify the frequency and explore potential mechanisms of sex ratio distortion. Strikingly, more than one-third of CC strains yield significantly sex-biased litters. These sex biases are not mediated by environmental effects and are moderately heritable. We conclude that the widespread sex ratio distortion in the CC manifests from multilocus permutations of selfish sex-linked elements and suppressors that are only recovered in the recombinant CC strains.

## INTRODUCTION

In species with single locus sex determination, Mendel’s rules of inheritance predict an equal number of males and females at birth. However, departures from this idealized expectation are common in nature. Many species can manipulate offspring sex ratios based on prevailing environmental conditions (Hamilton 1967; Nager *et al.* 1999; West and Sheldon 2002), including season (Drickamer 1990), maternal diet (Rosenfeld *et al.* 2003), and resource availability (Douhard 2017). Additionally, increasing maternal stress loads lead to skewed sex ratios in many mammals (Linklater 2007; Helle *et al.* 2008; Ideta *et al.* 2009; Ryan *et al.* 2012).

Beyond environmental influences, sex ratio biases can also arise from diverse genetic mechanisms. At one extreme, a segregating X-linked recessive lethal allele will be disproportionately associated with male lethality and distort the population sex ratio toward an excess of females. Although selection should rapidly eliminate such a hypothetical variant from the population, many mutations exhibit sex-specific phenotypic effects (Karp *et al.* 2017) and could, therefore, contribute to sex biases in live birth ratios (e.g., McNairn *et al.* 2019). In addition, prior work from diverse natural and laboratory model systems has demonstrated that sex-linked selfish elements can drive sex ratio distortion (SRD) by promoting the transmission of their resident chromosome at the expense of the other sex chromosome (Wood and Newton 1991; Seehausen *et al.* 1999; Cocquet *et al.* 2012; Unckless *et al.* 2015; Lindholm *et al.* 2016; Helleu *et al.* 2016; Zanders and Unckless 2019; Courret *et al.* 2019).

Under most circumstances, the presence of an X-linked (or Y-linked) selfish element will impose an intense selective pressure for the rapid emergence of a suppressor on the other sex chromosome, thereby restoring a balanced population sex ratio. Consequently, at any given point, a population is expected to be at or near sex ratio parity due to the action of counteracting drive and suppressor elements. However, because different populations evolve independent drive-suppressor systems, outcrossing can disrupt co-adapted systems of alleles to unmask the presence of cryptic sex ratio distorters. Indeed, SRD is frequently observed in inter-population (James and Jaenike 1990; Merçot *et al.* 1995), intersubspecific (Macholán *et al.* 2008; Phadnis and Orr 2009; Good *et al.* 2010; Cocquet *et al.* 2012; Kruger *et al.* 2019), and interspecific hybrids (Dermitzakis *et al.* 2000; Tao *et al.* 2001). SRD is also typically associated with reduced fertility (Phadnis and Orr 2009; Cocquet *et al.* 2010, 2012; Zanders and Unckless 2019), implying that selfish drive elements often impose a fitness cost.

The house mouse *(Mus musculus)* sex chromosomes are crucibles of historical genetic conflict between feuding drive elements on the X and Y. More than 90% of the house mouse Y chromosome is comprised of ampliconic genes with X-linked homologous partners that are theorized to reflect historical bouts of driver-suppressor evolution (Mueller *et al.* 2008; Soh *et al.* 2014). One such family includes the SYCP3-like gene family members, *Slx, Slxl1,* and *Sly*, which are present at upwards of 50-100 copies per genome (Scavetta and Tautz 2010; Morgan and Pardo-Manuel de Villena 2017). The Y-linked gene *Sly* and its X-linked homologs *Slx* and *Slxl1* are selfish drive elements that promote the transmission of their parent chromosome, although the molecular mechanisms by which this distortion is rendered are not fully understood. Knockdown or deletion of *Sly* results in over-transmission of the X-chromosome and female biased-litters (Conway *et al.* 1994; Cocquet *et al.* 2009). Conversely, knockdown or deletion of *Slx/Slxl1* yields male-biased litters (Cocquet *et al.* 2012; Kruger *et al.* 2019). Although there are striking differences in *Slx/Slxl1* and *Sly* copy number among house mouse subspecies, the ratio of X-to Y-linked copies within subspecies has been maintained in stochiometric balance over evolutionary time, presumably as a result of selection for balanced sex chromosome transmission (Scavetta and Tautz 2010; Morgan and Pardo-Manuel de Villena 2017). As expected, experimental crosses between subspecies and wild-caught intersubspecific F1 house mouse hybrids frequently sire sex-biased litters (Macholán *et al.* 2008; Good *et al.* 2010; Turner *et al.* 2012).

Regardless of whether it emerges from environmental or genetic causes, SRD imposes profound impacts on populations. Departure from a 1:1 sex ratio can influence the rate of population growth, modulate the degree of mate competition, and even modify life-history trajectories (Hamilton 1967; Le Galliard *et al.* 2005; Székely *et al.* 2014). In addition, SRD can cause relative levels of diversity and divergence across the sex chromosomes and autosomes to deviate from their theoretical expectations (Ellegren 2009; Wilson Sayres 2018). Further, given that many diseases differ in incidence between the sexes (Ober *et al.* 2008; Regitz-Zagrosek 2012), even subtle shifts in the sex ratio can lead to profound changes in the disease burden of a population.

Despite its critical importance for the conservation of biodiversity, population dynamics, and its role in shaping the genomic architecture of heterogametic vertebrate sex chromosomes, the underlying biological mechanisms of SRD often remain elusive. In cases of environmentally-induced SRD, we typically lack a comprehensive mechanistic understanding of how information gathered from the maternal environment is biochemically relayed to the reproductive track to manifest SRD (Krackow 1995; Navara 2013). Moreover, the genetic mechanisms that fuel intergenomic conflicts between the sex chromosomes and lead to SRD are currently understood at limited molecular resolution (Bravo Núñez *et al.* 2018; Courret *et al.* 2019; Kruger and Mueller 2021). On-going efforts to address these key knowledge gaps would be well-served by the availability of robust and reproducible animal models equipped with powerful genomic resources and tools for genetic engineering.

Toward this goal, we performed an exploratory analysis of the Collaborative Cross (CC) mouse population to describe the prevalence and define potential causes of SRD in a premiere mammalian model system (Churchill *et al.* 2004). The CC is an 8-way recombinant inbred panel of mice developed from eight genetically diverse parental strains: A/J, C57BL/6J, 129S1/SvImJ, NOD/LtJ, NZO/H1LtJ, CAST/EiJ, PWK/PhJ, and WSB/EiJ. Five of these founder strains (A/J, C57BL/6J, 129S1/SvImJ, NOD/LtJ, NZO/H1LtJ) are classical inbred mouse strains of predominantly *M. m. domesticus* ancestry (Yang *et al.* 2007). CAST/EiJ, PWK/PhJ, and WSB/EiJ are wild-derived inbred strains representing each of the three cardinal house mouse subspecies (*M. m. castaneus, M. m. musculus,* and *M. m. domesticus,* respectively). The contribution of genetic material from three divergent subspecies, each with variable *Slx/Slxl1* and *Sly* copy numbers, led us to specifically hypothesize considerable scope for genetic SRD in this multiparent mapping population.

We collated detailed breeding records from 58 genetically distinct CC strains maintained and distributed by The Jackson Laboratory to define patterns of SRD across this strain resource. We find that weak SRD is widespread in the CC. We integrate in-depth analyses of breeding records with genomic and phenotypic analyses to test the potential action of multiple mechanisms of SRD. Taken together, our findings underscore the untapped potential of the CC mouse mapping population to serve as a tool for dissecting the complex genetic mechanisms and environmental drivers of SRD.

## METHODS

### Compilation of Collaborative Cross breeding records

Breeding records were obtained for 58 Collaborative Cross strains maintained in The Jackson Laboratory’s Repository between January 2016 and July 2019. Eleven strains sired <100 pups surviving to wean age during this time frame and were excluded from further analyses. The data recorded for each live-born litter include: strain name, unique dam and sire identifiers, dam date of birth, sire date of birth, date mating established, litter birth date, litter size at birth, litter size at wean, and the number of weaned males and females. All CC breeding data are provided in **Table S1**. Strain-level summaries of these breeding data are provided in **Table S2**.

Breeding records for CC mice maintained at the University of North Carolina Chapel Hill (UNC) were obtained from the UNC Systems Genetics website (http://csbio.unc.edu/CCstatus/index.py?run=availableLines). These data are also made available in **Table S3**.

Breeding records for 43 F1 crosses between distinct CC strains (i.e., CC-RIX crosses) were kindly shared by colleagues at The Jackson Laboratory and are provided in **Table S4**.

For brevity, we exclude the laboratory code from CC strain names in figures and tables throughout this manuscript. Note that in all cases of such ambiguity, we are referencing CC lines maintained at the Jackson Laboratory.

### Estimating sex ratios and survival statistics

For most analyses, CC strain sex ratios were calculated as the proportion of females at wean. Sex ratios for CC lines maintained at UNC are presented in public data as the ratio of females to males at wean. Comparisons of CC strain sex ratios between the JAX and UNC breeding centers utilize JAX CC strain sex ratios calculated per this alternative definition.

Neonatal survival was approximated as the fraction of pups born to a given strain that survive to wean. We acknowledge that this estimate is potentially imprecise, as it is often difficult to accurately count pups at birth and pups that were cannibalized shortly after birth are likely missed in these tallies. Litter size at birth was used as a proxy for the *in utero* survival rate. Although litter size is shaped by a multitude of factors, strains with smaller litters may experience higher rates of embryonic lethality than strains with larger litter sizes, all else being equal.

To estimate survival during early *in utero* development and throughout the neonatal period, we devised an *ad hoc* metric that combined litter size at birth and survival to wean. Specifically, we computed the median litter size at birth and median birth-to-wean survival rate across all CC strains. For a particular focal CC strain, we then computed the difference between the strain-specific litter size and the overall CC population-wide median litter size. Similarly, we calculated the difference between the strain-specific survival rate and the overall median survival rate in the CC population. We then summed these two values into a single measure of aggregate strain-specific survival from conception to wean.

### Linear modeling and statistical analyses of temporal changes in SRD

The sex of each weaned pup was coded as a binomial indicator and modeled as a function of strain, birth month, and birth year using the *glm* function in RStudio (v. 1.3.1056). Post-hoc Wald tests were used to determine whether any independent variables provide a significant explanatory effect. R code to recapitulate these findings is available as a supplementary document (cc_srd_analysis.R) on FigShare.

### Analyses of maternal condition and male reproductive phenotypes

Maternal body mass and body fat percentage estimates for CC strains were obtained from the McMullan1 and McMullan3 datasets in the Mouse Phenome Database (**Table S5**; (Bogue *et al.* 2020)). We also accessed male reproductive phenotype datasets for CC (Lazear, Shorter3, Shorter4) and parental inbred strains (Odet1; (Odet *et al.* 2015)) via the Mouse Phenome Database (**Table S6**). Spearman Rank correlations and Mann-Whitney U-tests were used to assess relationships between phenotypes and SRD. R code to replicate analyses of maternal condition (maternalCondition.R) and reproductive phenotypes (ReproPhenotypeAnalysis.R) is available on FigShare.

### Broad-sense heritability estimation

We treated the sex ratio estimated from weaned pups born to individual CC mating units as independent, within-strain replicate phenotype measures. Mating units producing fewer than 30 pups over their breeding history were excluded due to the high uncertainty in calculated sex ratios. We then fit a one-way ANOVA model (sex ratio ~ strain) to estimate the broad-sense heritability *(H*^2^) of the sex ratio at wean using the interclass correlation method (Rutledge *et al.* 2014):

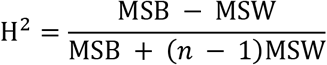

where MSB and MSW are the mean squares between and within strains, respectively. Despite being expressed as a proportion, the distribution of sex ratio values across the CC population is bell-shaped, and we confirmed that a square root transform of the data has no meaningful impact on the magnitude of our *H*^2^ estimate.

### QTL Mapping

Single QTL mapping was performed using the linear mixed model framework implemented in the R/qtl2 package (Broman *et al.* 2019). Genotype data at 110,054 unique genomic positions in each individual CC strain and the CC linkage map were accessed from Dr. Karl Broman’s GitHub page at https://raw.githubusercontent.com/rqtl/qtl2data/master/CC/cc.zip. Raw genotypes were converted to genotype probabilities by invoking the Carter-Falconer mapping function and assuming an error probability of 0.002. Relatedness among CC lines was specified via a kinship matrix using the leave-one-chromosome-out method. QTL significance thresholds were computed from 1000 permutations of the data.

We tested for significant effects of the Y and mitochondrial chromosomes on SRD using oneway ANOVA, treating the strain origin of the Y chromosome (or mitochondrial genome) as a factor.

R code to replicate QTL mapping and testing for Y and mitochondrial effects is available on FigShare (qtlMapping_sexRatio.R).

### Read mapping and Structural Variant Discovery

Whole genome sequences for the completed CC strains were previously released for community use (Srivastava *et al.* 2017; Shorter *et al.* 2019b). Fastq files for each CC sample were downloaded from the ENA archive under Project PRJEB14673. Fastq files for five CC lines (CC019, CC026, CC049, CC070, and CC076) were corrupted and we were unable to pursue genomic analyses with these strains.

Fastq files were first processed with *fastp* (version 0.20.1) to remove adaptor sequences and evaluate QC metrics (Chen *et al.* 2018). Reads were then mapped to the mm10 reference assembly using default parameter settings in *bwa mem* (version 0.7.17) (Li 2013). Optical duplicates were marked using the MarkDuplicatesSpark command within GATK (version 4.1.8.1; (Van der Auwera and O’Connor 2020)). Structural variant (SV) discovery was performed individually on each CC genome using *manta* (version 1.6.0; (Chen *et al.* 2016)). The resulting SV vcf files were reformatted using the python script *convertInversion.py* supplied with the manta distribution and subset to include only the non-ampliconic portion of the Y chromosome (chrY:1-6.664Mb). The resulting vcf files were then uploaded into the Integrative Genomics Viewer (IGV; version 2.8.0; (Robinson *et al.* 2011)) for manual inspection.

### Estimation of gene copy number for ampliconic sex-linked families

Depth of coverage was computed in 1kb non-overlapping sliding windows using *mosdepth* (version 1.18), ignoring read duplicates (Pedersen and Quinlan 2018). Coverage values were then corrected for GC biases using a custom R script (compute_GC_depth_correction.R; available on FigShare). Briefly, for each strain genome, we computed the average read depth across autosomal regions with identical GC content, excluding outlier windows with >2× and <0.333× the average autosomal coverage. We then used LOESS regression to fit a second-degree polynomial (span parameter = 0.7) to the data to model the empirical relationship between GC-content and average depth across the genome. The difference between the genome-wide average read depth and the predicted read depth for a given GC-content value corresponds to the average over- or under-representation of sequenced reads derived from regions in that GC-content bin. These values were used as correction factors to adjust the observed read depth in a given 1kb window based on its GC-content. GC-corrected read depths were then standardized by the average genome-wide coverage to convert to copy number estimates. Finally, data were compiled in bedGraph format for custom visualization in IGV. bedgraph format files are provided on FigShare.

To estimate the copy number of the X and Y-linked ampliconic genes *Slx/Slxl1, Sly, Sstx,* and *Ssty1/2,* we first identified the reference coordinates of all annotated paralogs from these genes using the Ensembl Paralogues feature. To discover any additional unannotated paralogs, we blated each annotated paralog sequence against the mm10 reference, retaining full-length hits with >90% sequence identity to the parent sequence. Genomic coordinates for all ampliconic genes are provided in **Table S7**. For each CC strain, we then summed the average read depth across all paralogs to obtain an overall estimate of gene family copy number.

### Copy number quantification by droplet digital PCR

*Slx*, *Slxl1,* and *Sly* copy number states were independently confirmed in a representative sample of six CC strains (CC032/GeniUncJ, CC011/UncJ, CC003/UncJ, CC004/TauUncJ, CC028/GeniUncJ, CC061/GeniUncJ) using droplet digital PCR (ddPCR). Genomic DNA was isolated from spleen tissue using a Qiagen DNAeasy kit following manufacturer recommendations. DNA was subsequently restriction digested with *HaeIII* for 60 minutes at 37C. DNA was then diluted 1:100 and 1:10 and combined with QX200 ddPCR EvaGreen Supermix (BioRad) and custom-designed primers targeting either *Slx/Slxl1, Sly,* or the diploid control *Rpp30* locus (**Table S8**) per vendor protocols. The PCR reaction mixture was then partitioned into droplets using a QX200 AutoDG Droplet Generator (BioRad). Emulsified reactions were cycled on a C1000 Touch Thermocycler (BioRad) according to the following program: 5 min initial denaturation at 95C; 40 cycles of 30s elongation at 95C, 30s of annealing at 55C, and 60s elongation at 72C; and a final 15 min incubation at 4C to stabilize fluorescent signals. Completed reactions were then held at 20C until removal from the thermocycler. Finally, reaction products were loaded into a QX200 Droplet reader and analyzed using QuantaSoft Analysis Pro Software (v. 1.0.596; BioRad). Copy number estimates were standardized to *Rpp30* and averaged across a minimum of two technical replicates at each DNA concentration. Final copy number estimates were then averaged across the two analyzed concentrations.

### Animal Husbandry and Use Statement

Collaborative Cross strains were obtained from the Jackson Laboratory’s Repository and housed in a low barrier room in accordance with an animal care protocol approved by The Jackson Laboratory’s Animal Care and Use Committee (Protocol # 17021). Mice were provided with food and water *ad libitum.* Sexually mature males were euthanized by exposure to CO2 at 10-14 weeks of age.

### Testis Histology

Whole testes from three CC032/GeniUncJ males were fixed in Bouin’s fixative overnight at 4C and then rinsed in a sequential ethanol series (25%, 50%, 3x 70% for 5 minutes each). Fixed tissues were then submitted to the Histopathology Sciences Service at The Jackson Laboratory for paraffin embedding, 5 μm cross-sectioning, and regressive staining with Mayer’s Hematoxylin and Eosin-Y. Slides were then scanned on a Hamamatsu NDP Nanozoomer at 40x magnification and analyzed using NDP.view2 software.

Two independent cross-sections from two of the three biological replicates were then scored for the following testis histology phenotypes: the total number of tubules per cross-section, total cross section area, the number of tubules with vacuoles, the number of tubules with eosinophilic cells (a proxy for cell death), and the cross-sectional area of 50 representative tubules. Testis cross-sections from the third CC032/GeniUncJ replicate exhibited an unusually high fraction (>50%) of vacuolized tubules harboring no post-meiotic cells (**Figure S1**). Inclusion of data from this sample vastly skewed the estimates from other replicates and was dropped from the analysis. Due to the limited number of surveyed replicates, we cannot exclude the possibility that this vacuolized seminiferous tubule phenotype affects a reproducible subset of CC032/GeniUncJ males, as previously reported for inbred WSB/EiJ males (Odet *et al.* 2015).

### Sperm whole chromosome painting

Sperm specimens were prepared for whole chromosome painting as described (Sarrate and Anton 2009; Dumont 2017). Sperm were passively isolated from the caudal epididymis in a drop of sterile PBS at room temperature, then concentrated by centrifugation and gradually resuspended in ~1mL of Carnoy’s fixative. Several drops of fixed sperm were then spread across a cleaned glass slide and allowed to air-dry. Slides were then rinsed in two successive washes of 2x SSC for 3 min each, dehydrated in a sequential ethanol series (70, 90, 100% for 2 minutes at each concentration), and air-dried. Next, slides were washed in dithiothreitol solution (5 mM 1,4-dithiothreitol, 1% Triton X-100, and 50 mM Tris) to decondense sperm DNA and rinsed again in two consecutive washes of 2x SSC, dehydrated in a sequential ethanol series (70, 90, 100%), and air-dried.

Sperm DNA was denatured in 70% formamide/2x SSC at 78C for 5 min. Slides were then processed through a sequential ethanol series (70%, 85%, 100% for 1 min each dilution) and air-dried. Simultaneously, Texas-Red labeled X chromosome and FITC-labeled Y chromosome probes (Cytocell) were denatured for 10 minutes at 80C per vendor recommendations. Sperm slides were then painted with a total volume of 10 μL of denatured probes, a cover slip was applied to the hybridized area, and sealed in place with rubber cement. Hybridization reactions were allowed to process for ~48 hr at 37C.

After removing coverslips, slides were washed in 0.4x SSC/0.3% NP-40 at 74C for 2 minutes, and 2xSSC/0.1% NP-40 at room temperature for 1 minute. After air-drying, slides were mounted in ProLong Gold antifade media with DAPI (Invitrogen) and cover slipped.

Approximately ~1400 painted sperm cells were imaged on a Leica DM6 B upright epifluorescent microscope equipped with GFP and Texas Red fluorescent filters and a cooled monochrome 2.8-megapixel digital camera. Images were post-processed and analyzed in the Fiji software package (Schindelin *et al.* 2012). Individual sperm were scored as carrying an X or Y chromosome based on fluorescent signal (**Table S9**). As no significant difference in the frequency of X-versus Y-bearing sperm was noted in an initial sample, we did not perform experiments on additional biological replicates or dye-swap probe combinations.

### Sex ratio distortion in the Diversity Outbred population

Breeding records for the 175 Diversity Outbred (DO) breeding lineages were obtained from the supplemental material of (Chesler *et al.* 2016) and are available in **Table S10**. These records detail the parentage of litters born over 17 continuous generations of outbreeding (G6-G22). To identify DO lineages siring sex-biased litters, we employed binomial tests to ask whether the sex ratio of weaned pups born to females (or males) from each DO breeding lineage deviated from the expected 1:1 ratio. An R script for reproducing this analysis is available on FigShare (do_srd_analysis.R).

### Data availability

The authors state that all data necessary for confirming the conclusions presented in this article are represented fully within the article and supplemental material. R scripts for reproducing analyses and figures are available on FigShare. VCF files with structural variant calls are available upon reasonable request from the corresponding author.

## RESULTS

### Widespread Sex Ratio Distortion in the Mouse Collaborative Cross

We collated breeding records from the Collaborative Cross mouse colony maintained in The Jackson Laboratory’s Repository (**Table S1**). These records summarize the breeding performance of 58 inbred CC strains organized into 3,890 independent mating units that produced 54,034 pups between January 2016 and May 2019 (median = 874 pups per strain). Eleven strains produced fewer than 100 pups during this time period and were excluded from further analysis. Remarkably, 18 of the remaining 47 CC lines (38%) sired progeny with a significant departure from the expected 1:1 sex ratio (uncorrected binomial *P* < 0.05; **Figure 1**). Of these 18 strains, 12 are significantly male-biased, whereas six produce an excess of females. This finding aligns with the significant, albeit slight, skew toward males in the aggregate CC population breeding records (21,469 males versus 20,439 females; two-sided binomial *P* = 4.99×10^-7^). The most significantly female-biased strain, CC065/UncJ, produces 67.4% females (n = 264; Binomial Test *P* = 1.54×10^-8^). CC032/GeniUncJ is the most male-biased strain, siring 68.6% males at wean (n = 927; *P* = 2.83×10^-30^).

**Figure 1.**
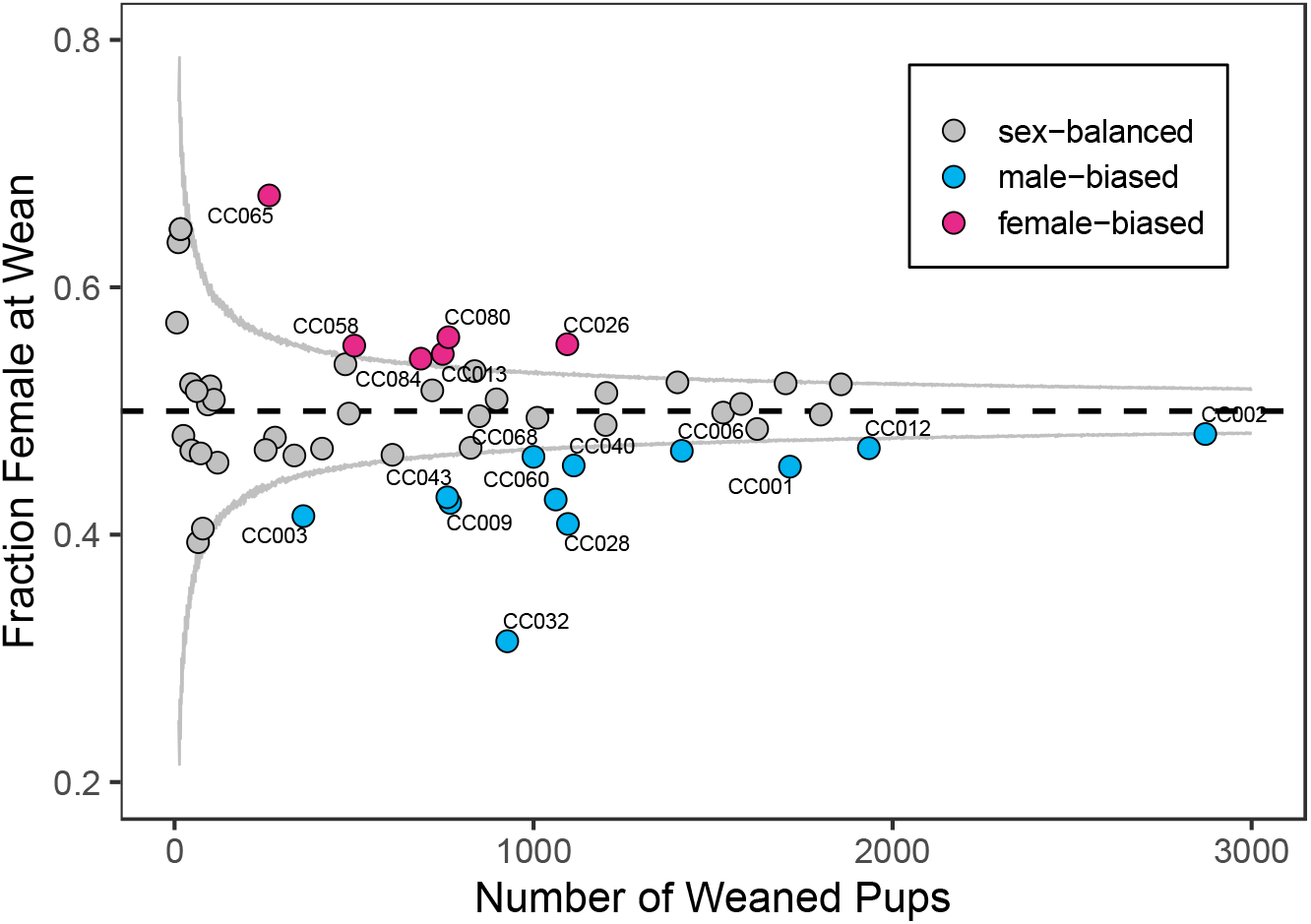
Sex ratio distortion in the realized Collaborative Cross population. Sex ratio is expressed as the fraction of females at wean. The gray curved lines denote the level of noise about the Mendelian expectation of 0.5 (dashed horizontal black line) that can be expected for a given sample size due to binomial sampling error. Strains producing significantly male- and female-biased litters are color-coded blue and red, respectively.

Statistical power to detect a significant departure from the expected Mendelian sex ratio is a function of sample size. Many CC strains produced a modest number of pups over the survey period, limiting our ability to detect weak SRD. Considering only those strains with >500 pups (corresponding to ~60% power to detect a 45%:55% skew in the sex ratio; **Figure S2**), the percentage of strains with significant SRD increases to 50%. We conclude that mild SRD is pervasive in the CC reference mapping population, with a few strains showing extreme biases in offspring sex ratios.

### Testing the Stability of SRD to Environmental Influences

Seasonal fluctuations in external temperatures can influence offspring sex in captive laboratory house mouse populations (Drickamer 1990). To test for seasonal and larger-term temporal effects on SRD in the CC, we modeled the sex of each weaned pup as a binomial outcome of strain identity, birth month, and birth year. Neither birth month nor birth year provide significant predictive power in this model (birth month, Wald Test *P* = 0.74; birth year, Wald Test *P* = 0.17). These findings are recapitulated on a per strain basis: there is no evidence for variation in sex ratio from month-to-month within strains (Fisher’s Exact Test, *P* > 0.05; **Table S11**). CC028/GeniUncJ and CC084/TauJ show slight variation in sex ratio from year-to-year (**Figure S3**; Fisher’s Exact Test; *P*_CC028_=0.0365 and *P*_CC084_=0.0500), although these effects are modest and do not remain significant after correcting for multiple testing.

To understand whether housing environment influences SRD in the CC, we next assessed the concordance of JAX CC strain sex ratios with those of their sister-strain counterparts maintained at an independent mouse facility at University of North Carolina (UNC) Chapel-Hill. Overall, there is excellent concordance of the strain sex ratios between these two locations (Spearman’s Rho = 0.760, *P* = 1.97×10^-8^; **Figure 2**). Notably, CC032 and CC065 are the most strongly male- and female-biased strains, respectively, regardless of facility. Despite this overall agreement, there are minor exceptions. For example, two strains that produce slightly male-biased strains at JAX – CC060/UncJ and CC002/UncJ – fail to produce male-biased litters at UNC. These slight discrepancies are likely attributable to binomial sampling error, rather than the accumulation of independent mutations with effects on offspring sex ratio in the JAX and/or UNC colonies *(i.e.,* strain drift) or strain-by-environment interactions.

**Figure 2.**
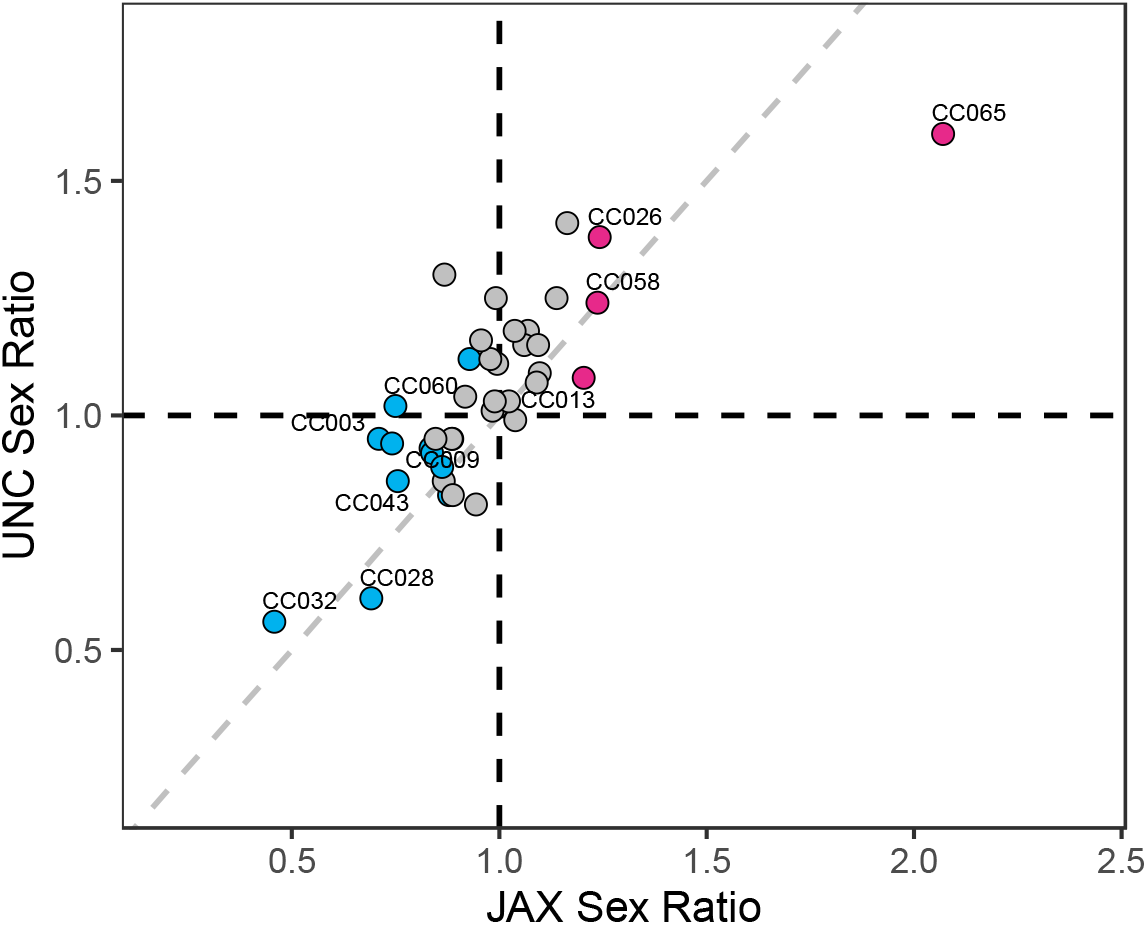
Sex ratios of CC strains maintained at JAX and UNC. Sex ratios on both axes are expressed as the number of females to the number of males at wean. Strains that are significantly female- and male-biased at JAX are color-coded red and blue, respectively. The dashed gray diagonal line represents y=x.

### Sex ratio is not influenced by maternal effects or breeding performance

Maternal condition has been shown to modulate live-birth sex ratios in many organisms (Nager *et al.* 1999; Love *et al.* 2005). Using publicly available CC mouse phenotypes (Bogue *et al.* 2020; **Table S5**), we examined the relationship between sex ratio and two proxies of overall maternal health: adult female body mass and body fat percentage. Sex-biased strains do not have significantly different body mass or body fat percentages compared to sex-balanced strains (Mann-Whitney U-Test; *P* > 0.05; **Figure S4**). Similarly, there is no significant difference in body mass between female-biased strains and sex-balanced strains, or between male-biased strains and sex-balanced strains (Mann-Whitney U-Test; *P* > 0.05). Although we find no significant link between these two metrics of overall maternal condition and offspring sex ratio, we acknowledge that female-specific estimates of condition, rather than the strain-wide estimates employed here, are most appropriate for rigorously testing this possible explanation for SRD.

We next considered the possibility that random, non-genetic maternal effects influence offspring sex in the CC. In the majority of CC strains, offspring sex does not vary from dam-to-dam within a strain (Fisher’s Exact Test; *P*>0.05; **Table S12**). We do observe slight fluctuations in sex ratio across breeding dams in CC004/TauUncJ (*P* = 0.013), CC061/GeniUncJ (*P* = 0.008), and CC068/TauUncJ (*P* = 0.033), although these effects are not significant after accounting for multiple testing (**Figure S5**; 42 tested strains, Bonferroni adjusted *P* = 0.0012).

Prior work has uncovered significant effects of parental age and litter parity number on mammalian sex ratios (Huck *et al.* 1988). Parental age at litter birth, litter size, and the number of litters born to each mating unit vary among CC strains (**Table S2**), prompting us to explore whether these variables contribute to the observed SRD. We modeled the sex of each weaned pup as a binomial outcome of strain identity and either dam or sire age. Parental age does not offer significant explanatory power in this model *(P* > 0.05). Similarly, litter size and litter number do not impact estimated sex ratios (*P* > 0.05).

### Sex ratio distortion is independent of maternal genotype

Different maternal genotypes provide distinct uterine environments for early development and could modify sex ratios in crosses involving sires from a common strain. We leveraged breeding data from crosses between distinct CC strains *(i.e.,* CC-RIX crosses) carried out by colleagues at The Jackson Laboratory to address whether maternal genotype influences CC sex ratios. In total, we surveyed data from 43 CC-RIX crosses profiling 24 different CC strains as sires (**Table S4**). If sex ratio is strictly determined by the paternal transmission of X-versus Y-bearing sperm, then the sex ratios of litters sired by males from different CC strains should be independent of dam genotype. For each of the 24 CC sire strains included in this CC-RIX dataset, we asked whether offspring sex varies as a function of dam strain identity. Although small sample sizes limit our power (range: 13-103 progeny per CC-RIX cross; median = 31), we find no consistent evidence for maternal genotype-dependence of sex ratios in the majority of the CC-RIX strains tested (Fisher’s Exact Test *P* > 0.05; **Table S13; Figures S6, S7**). Three CC strains are exceptions, with marginal maternal genotype dependence on offspring sex ratios: CC027/GeniUncJ (*P* = 5.2×10^-4^), CC042/GeniUncJ (*P* = 0.038), and CC060/UncJ (*P* = 0.010) (**Figure S6**). Notably, CC027/GeniUncJ males, when mated to CC027/GeniUncJ or CC011/UncJ females, produced sex-balanced litters, but yield female-biased litters in crosses to CC037/TauUncJ dams and male-biased litters in crosses to CC002/UncJ dams (**Figure S6B**). Similarly, CC042/GeniUncJ males produce litters with a slight female bias in crosses with either CC042/GeniUncJ or CC001/UncJ dams, and male-biased progeny in crosses to CC005/TauUncJ females (**Figure S6A**). However, we caution that only the maternal effects in CC027/GeniUncJ remain significant after correction for multiple tests. Overall, these findings are in broad agreement with the absence of significant maternal genotype effects on sex ratios in the eight parental founder strains of the CC (Shorter *et al.* 2019a).

In summary, we find no evidence that season, housing environment, dam identity, parental age, litter number, litter size, maternal condition, or maternal genotype systematically influence sex ratios across the CC strains. Based on these findings, we conclude that the sex ratio of a given CC strain is likely an intrinsic, biological property of that strain, rather than a plastic response to environmental factors or mediated via parental effects.

### Evaluating sex differences in survival as a potential mechanism of sex ratio distortion

SRD can arise along a continuum of developmental timepoints ranging from differences in the viability or fertilization efficiency of X-versus Y-bearing sperm to sex differences in post-birth survival. If SRD stems from sex differences in survival during *in utero* development, more extremely sex-biased strains should yield smaller litters. In contrast to this expectation, there is no correlation between litter size and the absolute deviation from sex ratio parity in the CC population (**Figure 3A**; Spearman’s Rho = −0.036, *P* = 0.819; analysis restricted to breeding pairs only, to the exclusion of breeding trios and harem breeding units). Indeed, several sex-biased strains – including CC013/GeniUncJ, CC060/UncJ, CC006/TauUncJ, and CC001/UncJ – are among the most fecund CC lines. Similarly, sex differences in survival from birth to wean are not correlated with SRD (**Figure 3B**; Spearman’s Rho = −0.0817, *P* = 0.601).

**Figure 3.**
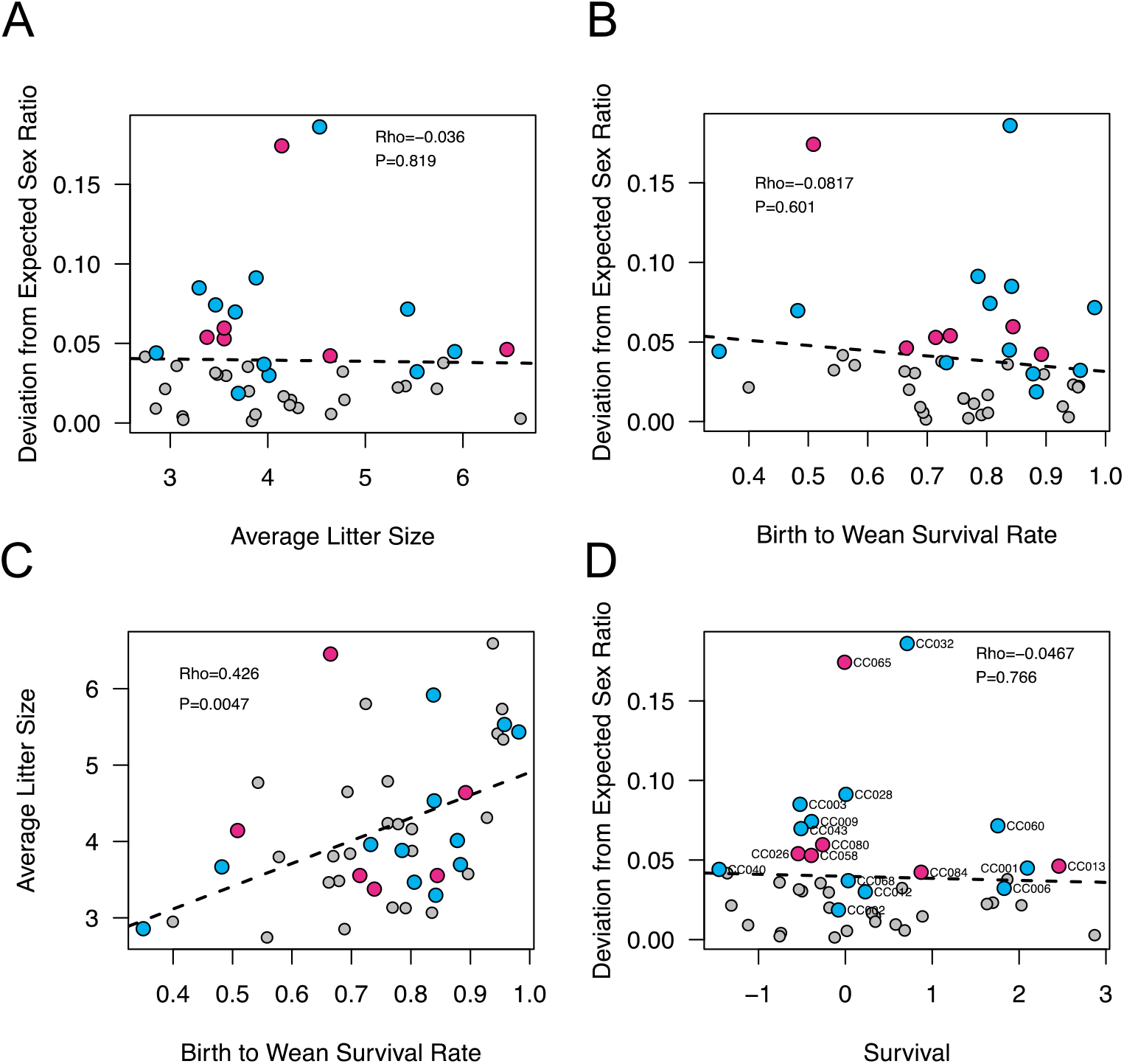
Correlations between neonatal survival, litter size, and sex ratio distortion. Sex ratio distortion is not significantly correlated with average litter size (A) or birth to wean survival rate (B). Average litter size at birth and survival to wean are positively correlated (C). The strength of sex ratio distortion is not correlated with aggregated *in utero* and birth-to-wean survival. Points corresponding to strains with significant female- and male-bias are color-coded red and blue, respectively.

Mechanisms that contribute to increased rates of strain death *in utero* may also lead to increased death rates in neonates. Consistent with this possibility, there is a significant positive correlation between litter size and birth-to-wean survival rate; strains with larger litters at birth have lower neonatal death rates (**Figure 3C**; Spearman’s Rho = 0.426, *P* = 0.005). We combined these two measures of survival during pre- and post-natal development into a single statistic that summarizes strain variation in survival from conception to wean (see Methods). Again, we find no significant correlation between this composite survival statistic and the magnitude of SRD (**Figure 3D**; Spearman’s Rho = −0.0467, *P* = 0.766).

The absence of an overall association between survival in early development and SRD suggests that mortality in early life does not provide a simple, unifying explanation for SRD in the CC. However, it is noteworthy that two of the 18 significantly sex-biased strains are among the 20% of CC strains with the lowest survival rates (CC026/GeniUncJ and CC040/TauUncJ; **Figure 3D**). Sex differences in early development may contribute to SRD in certain strains, and these findings motivate further work to dissect the developmental mechanisms of potential sexspecific mortality in these lines. Nonetheless, survival differences are unlikely to explain SRD in the majority of sex-biased strains.

### No Evidence for Single Locus Mechanisms of Sex Ratio Distortion

Our extensive analyses of possible non-genetic explanations for sex ratio variation in the CC turned up no compelling explanations, suggesting that sex ratio variation likely carries a genetic basis. We utilized sex ratio estimates from independent breeding units within each CC strain as biological replicates to compute the relative proportion of variation in the sex ratio that is due to within versus between strain differences *(i.e.,* broad sense heritability, *H^2^;* see Methods). Despite considerable binomial noise in sex ratio estimates per breeding unit, the sex ratio is modestly heritable in the CC *(H^2^* = 0.263).

To attempt to map genomic loci contributing to this heritable variation in SRD, we carried out a genome-wide QTL scan in the CC. No autosomal or X-linked loci reached the genome-wide threshold for significance (**Figure 4A**). Similarly, we find no effect of the Y chromosome or mitochondrial haplotype on SRD (one-way ANOVA, *P* > 0.05; **Figures 4B and 4C**). Although the small number of CC strains severely limits mapping power, we conclude that very large-effect, single-locus modifiers of SRD are not likely segregating in the CC population.

**Figure 4.**
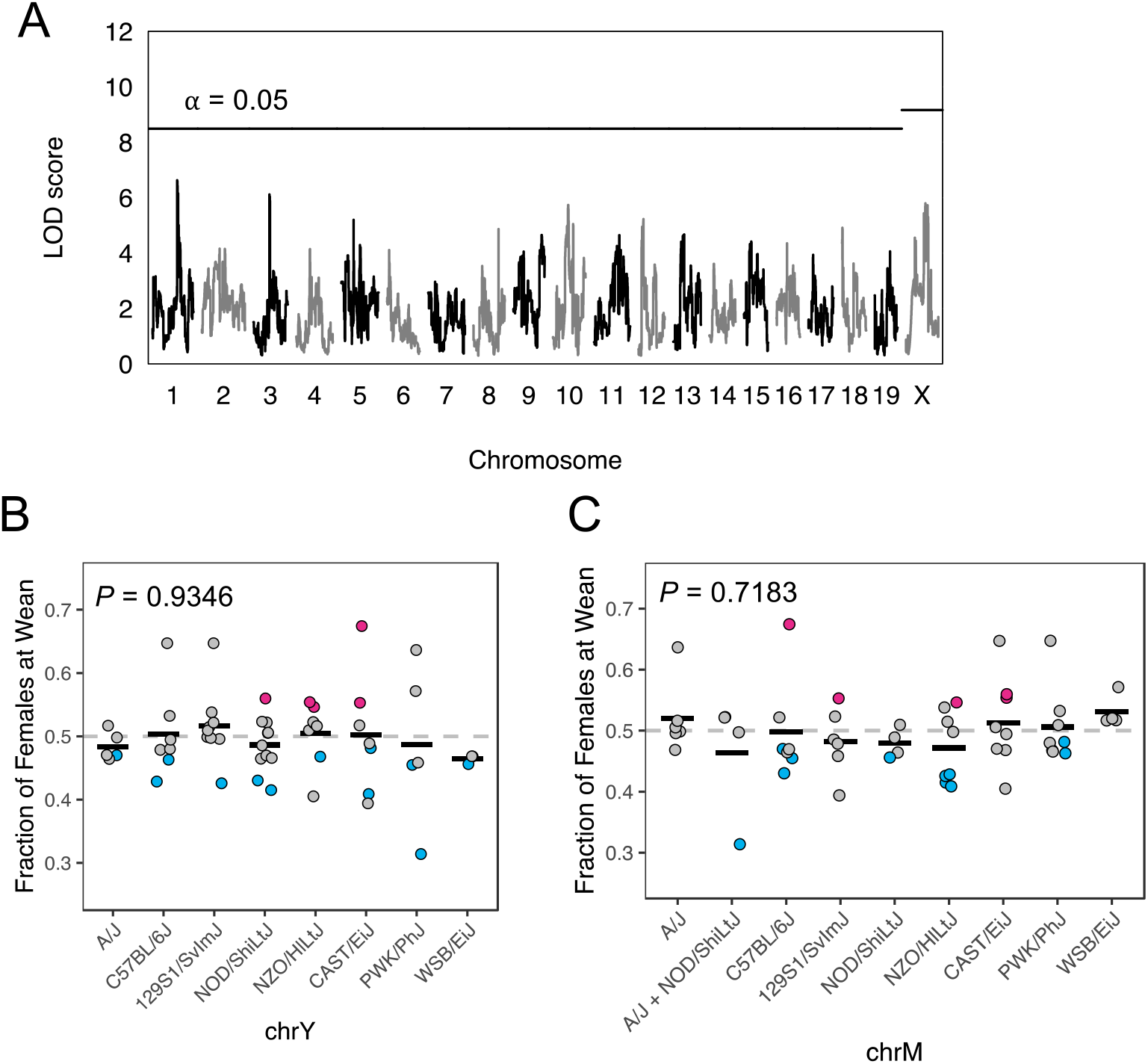
Mapping single locus modifiers of SRD in the CC. (A) Genome wide scan for loci influencing SRD in 49 CC strains. Horizontal lines correspond to the genome-wide permutation-derived thresholds for the autosomes and X chromosome. Fraction of females at wean for CC strains partitioned by the parental strain origin of the Y chromosome (B) and mitochondrial (C) haplotype. Strains with significantly male- and female-biased sex ratios are color-coded blue and red, respectively.

This conclusion is bolstered by the observation that sex ratios in seven of the eight inbred CC founder strains do not deviate from the expected 1:1 ratio of males:females (Shorter *et al.* 2019a). Founder strain 129S1/SvImJ produces a slight bias toward females (52.5%), but this distortion is mild compared to that observed in the most extreme CC strains. In conjunction with the overall absence of evidence for environmental or parental effects on SRD, these findings raise the possibility that multilocus allelic combinations only uncovered in recombinant CC genomes contribute to the widespread SRD in this multiparent mapping population. Unfortunately, the modest number of realized CC strains effectively bars the application of unbiased pairwise scans for such interacting loci.

### Structural mutations at sex determination genes are unlikely to mediate SRD in the CC

Structural mutations encompassing key sex determination genes can lead to disparities between phenotypic sex and sex chromosome genotype and could, potentially, manifest as SRD. In most mammals, including house mice, sex is determined by the expression of a *Y*-linked gene, *Sry,* in the undifferentiated gonad. Deletion or translocation of *Sry* from chrY represent established genetic mechanisms for sex reversal in mammals (McElreavy *et al.* 1992; Goodfellow and Lovell-Badge).

To explore the possibility that phenotypic sex is not a reliable indicator of sex chromosome transmission in the CC, we scanned *CC* whole genome sequences (Srivastava *et al.* 2017) for read mapping signatures consistent with structural mutations spanning *Sry* (see Methods). We uncovered no evidence for translocations or deletions encompassing the *Sry* locus in any CC strains. However, unexpectedly, we find that nearly all strains with the NOD/ShiLtJ Y chromosome carry a ~200kb duplication spanning the complete *Sry* coding region (**Figure S8**). CC003/UncJ is a single, notable exception: despite carrying a NOD/ShiLtJ Y chromosome, *Sry* is present as a single copy gene. Invoking parsimony, we conclude that a deletion of the duplicate *Sry* copy likely occurred during inbreeding of CC003/UncJ. A recent analysis of SNP array data in diverse mice identified a larger, 2.9 Mb *Sry*-spanning duplication in C3H/HeJ mice (Sigmon *et al.* 2020). The discovery of two independent *Sry*-spanning duplications in the classical inbred strains and a putative *de novo* deletion in CC003/UncJ suggests that this locus is inherently predisposed to recurrent genomic rearrangements and motivates further investigation into structural genetic diversity at this critical developmental regulator in house mice.

SRY activates the transcription of a second gene, *Sox9,* which in turn induces the testis developmental program. Duplication (deletion) of *Sox9* and/or its upstream regulatory elements can lead to constitutively high (low) levels of *Sox9* expression, providing a second mechanism for sex reversal in mammals (Foster *et al.* 1994; Gonen *et al.* 2018). Interestingly, while we find no evidence for duplication of *Sox9* itself, we observe a ~1kb duplication and a ~2kb deletion within the distal upstream *Sox9* regulatory region that are specific to animals carrying the NOD/ShiLtJ haplotype in these regions (**Figure S9**). These structural mutations do not span any annotated regulatory elements in the mm10 reference genome, but it is tempting to speculate that one or both may function to maintain native *Sox9* expression levels in the face of potentially increased SRY dosage driven by the *Sry*-duplication present in this strain.

It is unlikely that the NOD/ShiLtJ-specific SVs documented here are associated with an appreciable rate of sex reversal. NOD/ShiLtJ has served as a prominent mouse model of autoimmune disorders for more than 40 years, with no cases of sex reversal documented in this strain or in crosses involving this strain (including the CC), to our knowledge. In addition, *Sry* duplications are relatively common in rodent systems, and are not generally associated with sex reversal (Nagamine 1994; Lundrigan and Tucker 1997; Bullejos *et al.* 1999). Finally, half of the sex-biased CC strains do not carry the NOD/ShiLtJ haplotype at either *Sox9* or *Sry* (**Table S14**), necessarily assigning causality of SRD to other mechanisms. In summary, although we find novel structural rearrangements spanning the *Sry* sex determination gene and within the putative regulatory region of its upstream signaling target, *Sox9* (**Figures S8** and **S9**), these mutations seem unlikely, at face-value, to induce high rates of sex reversal and contribute to the widespread SRD in the CC population.

### Genetic Conflict mediated by Sex-linked Ampliconic Genes May Drive Sex Ratio Distortion in a Subset of CC Strains

The Y-linked ampliconic gene *Sly* and its X-linked counterparts, *Slx* and *Slxl1,* are embroiled in a genetic conflict over sex chromosome transmission during post-meiotic spermatogenesis (Cocquet *et al.* 2012; Good 2012). Given that the CC founder strains include representatives from three cardinal house mouse subspecies differing in their absolute *Slx/Slxl1* and *Sly* copy numbers (Morgan and Pardo-Manuel de Villena 2017), we hypothesized that *Slx/Slxl1* – *Sly* mediated conflict may underlie the pervasive pattern of SRD in this mapping population.

To address this possibility, we used publicly available whole genome sequences to estimate the relative genomic copy number of these ampliconic genes in each realized CC strain (**Table S15;** (Srivastava *et al.* 2017; Shorter *et al.* 2019b)). *Slx/Slxl1* copy number varies approximately 2.5-fold across strains (range: 24-64 copies), with only 5 strains exhibiting estimated haploid copy number states that fall outside the range delimited by the parental genomes (parental range: 28-60 haploid copies; **Figure S10A**). Across the CC population, *Sly* copy number ranges from 54-168. CC founder whole genome sequences were generated from female samples, barring comparisons of *Sly* copy number status in the parental inbred and realized CC strains.

Overall, we find no correlation between the fraction of females at wean and *Slx/Slxl1:Sly* copy number ratio (Spearman’s Rho = 0.0968, *P* = 0.490; **Figure 5A**). We validated the genomic read-depth *Slx/Slxl1:Sly* estimated copy number (CN) ratios using ddPCR assays in a representative subset of CC strains (**Table S16**). There is excellent qualitative alignment between these two orthogonal methods for copy number estimation (**Figure 5B**; Spearman’s Rho = 0.943, *P* = 0.0167), suggesting that the absence of a relationship between the *Slx/Slxl1:Sly* ratio and SRD is unlikely due to technical errors.

**Figure 5.**
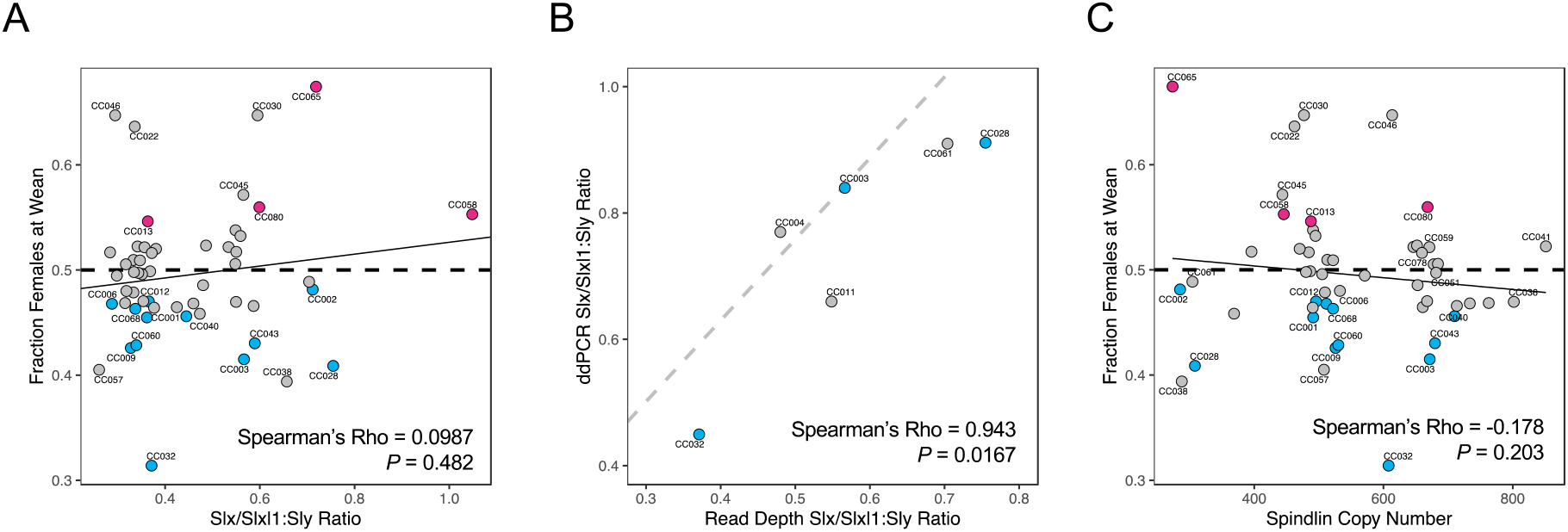
Relationship between copy number at ampliconic sex-linked genes and SRD. (A) Correlation between the fraction of females at wean and the ratio of Slx/Slxl1:Sly gene copy number. (B) Estimated copy numbers from genomic read depth and ddPCR are positively correlated for members of the SYCP3-like gene family. (C) Correlation between the fraction of females at wean and the copy number ratio of X:Y-linked genes in the spindlin gene family *(Sstx, Ssty1/2).* Points corresponding to strains with significantly male- and female-biased sex ratios are color-coded blue and red, respectively.

*Slx* and *Slxl1* are both neo-functionalized copies of SYCP3, but have rapidly diverged from each other, exhibiting just 61% protein identity (Kruger *et al.* 2019). Recent experimental work suggests that drive potential may be restricted to *Slxl1:* knockout of *Slxl1,* but not *Slx*, is associated with a shift toward male-biased litters (Kruger *et al.* 2019). We find that the estimated copy number of *Slxl1,* but not *Slx*, is positively correlated with SRD in the CC *(Slxl1:* Spearman’s Rho: 0.334, *P* = 0.0146; *Slx*: Spearman’s Rho = 0.227, *P* = 0.102). However, the ratio of *Slxl1: Sly* copy number carries no predictive association with SRD in this mapping population (Spearman’s Rho = 0.105, *P* = 0.455), in contrast to expectations.

Despite the lack of an overall association between the *Slx/Slxl1:Sly* copy number ratio and sex ratio distortion across CC strains, many sex biased strains do exhibit extreme amplicon copy number ratios. In particular, CC006/TauUncJ has a relative excess of *Sly* copies relative to *Slx/Slxl1,* consistent with the male bias in this strain. CC065/UncJ and CC058/UncJ, two female-biased strains, exhibit a relative excess of *Slx*/*Slxl1* compared to *Sly*, consistent with the over-transmission of the X chromosome. However, there are clear exceptions to expected trends. CC013/GeniUncJ has a moderately low *Slx/Slxl1:Sly* ratio, yet this strain is female-biased. CC002/UncJ and CC028/GeniUncJ have high *Slx/Slxl1:Sly* ratios, in contrast to predictions given the male sex bias observed in these strains (**Figure 5A**).

Recent molecular evidence suggests that SLX/SLXL1 and SLY compete for binding to SSTY1/2 and SPIN1, members of the spindlin gene family, to regulate expression at a large number of genes expressed during spermatogenesis (Comptour *et al.* 2014; Kruger *et al.* 2019; Moretti *et al.* 2020). SLY binding to SSTY1/2 or SPIN1 at gene promotors triggers the recruitment of the SMRT/Ncor complex, repressing gene expression. In contrast, SLX and SLXL1 do not interact with the SMRT/Ncor complex, and their association with SSTY1/2 at gene promoters leads to the upregulation of target genes (Moretti *et al.* 2020). *Ssty1/2* are Y-linked ampliconic genes, whereas the mouse X-chromosome harbors several *Spin1* gene clusters. The antagonistic interactions of *Ssty/Spin1* with *Slx/Slxl1* and *Sly* prompted us to explore whether copy number status at spindlin genes may factor into the complexity of SRD in the CC.

*Spin1* and *Ssty1/2* copy numbers span a 4.6- and 3.14-fold range in the CC population, respectively *(Spin1* range: 50-232 copies; *Ssty1/2* range: 223-698 copies; **Figure S10B**). We observe no significant relationship between the combined *Spin1* and *Ssty1/2* copy number and sex ratio (**Figure 5C**; Spearman’s Rho = −0.178, *P* = 0.203). Based on the known interactions between spindlins and members of the *Sycp3*-like family, we reasoned that the ratios of *Slx/Slxl1* to spindlin CN and *Sly* to spindlin CN might be correlated with the degree of SRD. These predictions are not upheld (Spearman rank correlation *P* > 0.05; **Figure S11**).

Overall, our findings uncover no global relationship between the copy number state of genes in the *Sycp3*-like and spindlin gene families with SRD in the CC. However, the copy number state of several CC lines accords with expectations under current models of SLX/SLXL1-SLY genetic conflict, and we speculate that this established drive system may contribute to SRD in an appreciable number of CC strains. Further, our genomic estimates of copy number for these ampliconic genes may not accurately estimate the number of transcriptionally active genes in each family. It is possible that a more widespread relationship between the copy number state of these ampliconic genes and SRD is concealed by the inclusion of large numbers of nonexpressed pseudogenes in our copy number tallies. Lastly, we cannot rule out the likely possibility that complex protein interactions between SLX/SLX1, SLY, spindlins and potentially other ampliconic spermatid-expressed gene families contribute to the SRD documented in the CC (Kruger *et al.* 2019; Moretti *et al.* 2020).

### Many sex-biased CC strains exhibit reduced male fertility

SRD is frequently associated with reduced fertility in experimental crosses (Phadnis and Orr 2009; Cocquet *et al.* 2010; Meiklejohn *et al.* 2018; Zanders and Unckless 2019; Kruger *et al.* 2019), a trend that may emerge from the differential death, motility, or fertilization capacity of sperm bearing one sex chromosome relative to the other. It is widely acknowledged that many CC lines are poor breeders, and it appears that in most cases, reproductive output is constrained by male fertility (Shorter *et al.* 2017). While there is no significant relationship between average litter size and SRD among CC strains (**Figure 3A**), we sought to assess the relationship between SRD and more precise measures of male fertility.

We accessed publicly available reproductive phenotype datasets for several CC strains from the Mouse Phenome Database (Bogue *et al.* 2018; **Table S6**). Although the limited number of phenotyped CC strains effectively bars a rigorous statistical analysis, many sex-biased strains do appear to have reduced fertility relative to sex-balanced strains. Sex-biased CC strains tend to have lower testis weights than non-sex-biased CC strains, although this association is not statistically significant (**Figure 6A**; Spearman’s Rho = −0.382, P = 0.248). Similarly, despite the lack of significant population-wide statistical correlations, several sex-biased strains – including CC028/GeniUncJ, CC032/GeniUncJ, and CC040/TauUncJ – exhibit low fractions of motile sperm and low sperm density compared to strains that yield sex-balanced litters (**Figure 6B,C**). We conclude that many sex-biased strains exhibit phenotypic signatures of reduced male fertility.

**Figure 6.**
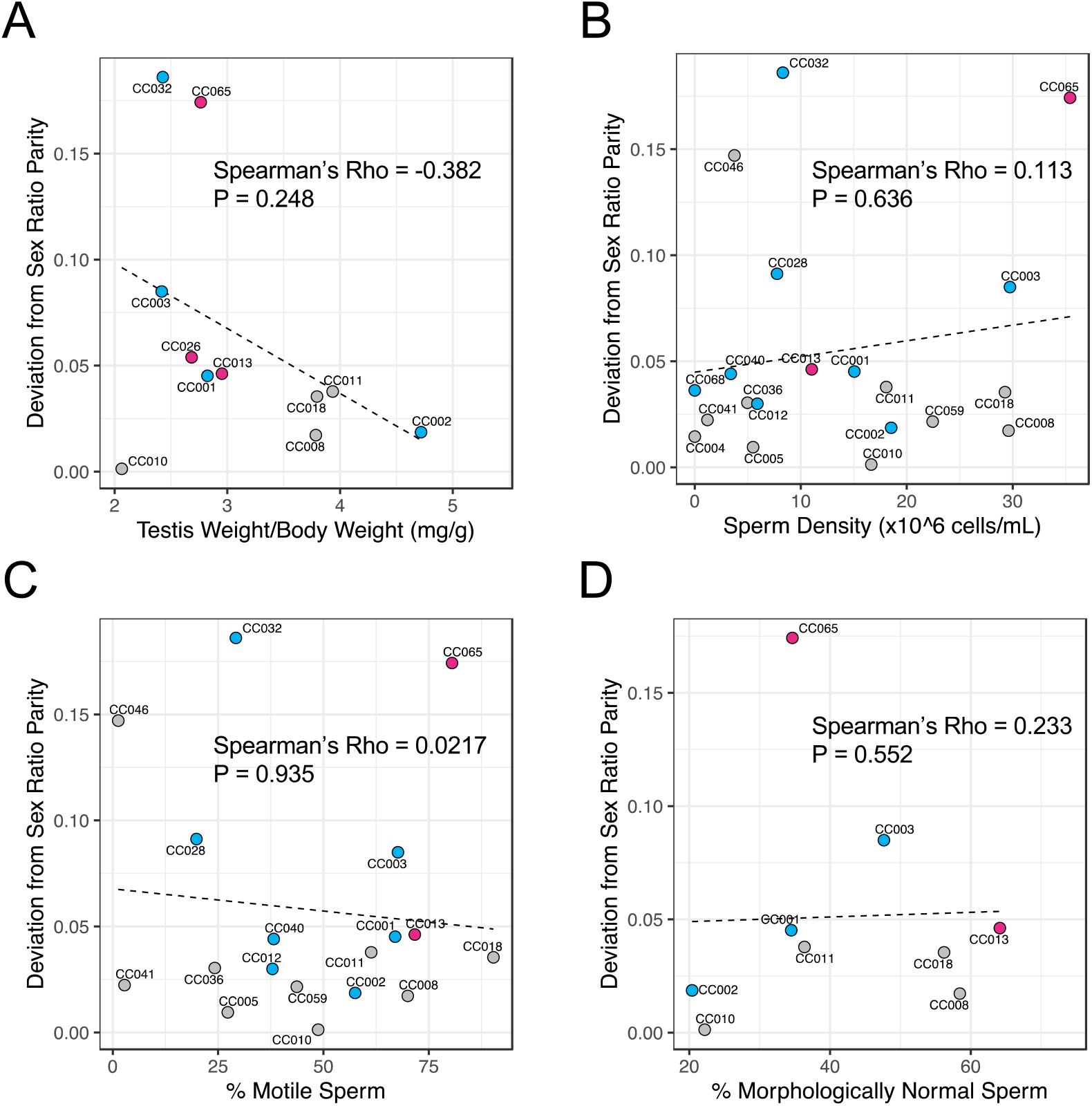
Correlations between the magnitude of sex ratio distortion and (A) testis weight standardized by body weight, (B) sperm density, (C) percentage of motile sperm, and (D) the percentage of morphologically normal sperm. Dashed black lines trend lines were derived from simple linear regression (y~x).

### Mechanisms of SRD in CC032/GeniUncJ

CC032/GeniUncJ (hereafter, CC032) is the most extremely sex-biased CC strain, with only one female out of every ~3 weaned pups. To our knowledge, SRD in this strain represents the strongest reported departure from Mendelian expectations in a mammal. Our analyses of strain breeding records indicate that SRD in CC032 cannot be entirely mediated by sex differences in survival. CC032 has moderately-sized litters (4.2 pups/litter; 73^rd^ percentile among CC strains) and intermediate rates of neonatal mortality (20.6% mortality from birth to wean; 42^nd^ percentile among CC strains). Remarkably, even if all live-born CC032 pups that did not survive to wean were female, this strain would still be significantly male-biased (**Table S2**; Binomial test *P* = 0.002565).

Relative to other CC lines, CC032 males have low average testis weights (**Figure 6A**), low sperm density (**Figure 6B**), and reduced motility (**Figure 6C**). Histological analysis of testis cross-sections reveal that CC032 males also have smaller seminiferous tubules than the majority of the CC founder strains (**Figure 7A**) and a higher fraction of tubules with vacuoles, (**Figures 7B-D**).

**Figure 7.**
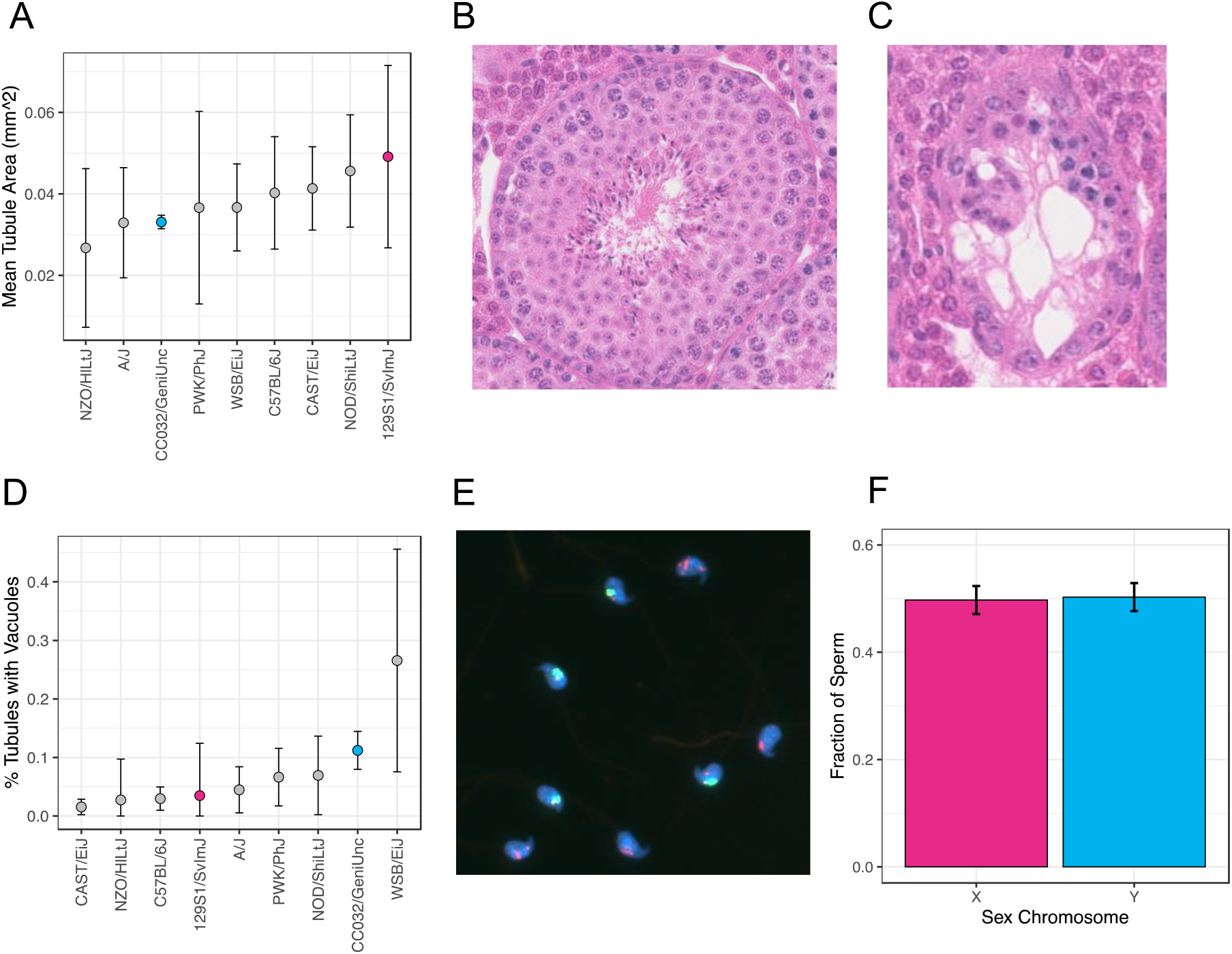
Reproductive phenotypes in CC032/GeniUncJ and the 8 CC founder strains. (A) Mean tubule area in mm^2^ (+/- 1 standard deviation). (B) Representative stage VII/VIII tubule cross section from CC032. (C) CC032 harbors a high fraction of tubules with large vacuoles. (D) Percentage of tubules with vacuoles (+/- 1 standard deviation) for each of the 8 CC founders and CC032. (E) Representative image of CC032 sperm hybridized with fluorescent paint probes against chrY (green) and chrX (red). Y-bearing sperm carry a slight signal from chrX due to shared X/Y homology across the pseudoautosomal region. (F) Fraction of X- and Y-bearing sperm in CC032 from X and Y chromosome painting.

Based on these phenotypic findings, we reasoned that targeted killing of X-bearing germ cells could be a plausible explanation for the observed male bias in CC032. We used whole chromosome painting to assess sex chromosome representation in mature sperm from this strain. We observe equal numbers of X- and Y-bearing sperm (49.7% chrX-bearing sperm; Binomial *P* = 0.854; **Figures 7E** and **7F**), dismissing this explanation for SRD.

CC032 harbors a PWK Y-chromosome with a high *Sly* copy number and an X chromosome bearing contributions from laboratory strains with *M. m. domesticus* ancestry *(e.g.,* moderate *Slx*/*Slxl1* copy number). As a consequence of this sex chromosome haplotype structure, the ratio of *Slx/Slxl1* to *Sly* in CC032 is lower than the CC-wide average (**Figure 5A**). The observed *Slx/Slxl1:Sly* ratio in this strain is consistent with the observed male-bias. However, the strength of SRD in CC032 exceeds that observed in both *Slx*/*Slxl1* knockdown and knockout mice (Cocquet *et al.* 2010; Kruger *et al.* 2019), suggesting the complementary action of other mechanisms that exacerbate SRD on this genetic background.

We scanned the genome of CC032 for potential structural mutations in other sex-linked ampliconic genes that could, conceivably, amplify the strength of *Slx/Slxl1*: *Sly*-mediated SRD. Strikingly, the CC032 genome shows a pronounced enrichment of reads mapping to chrXA.1 (chrX:3-6Mb; **Figure S12**) and chrXqA3 (chrX:30.5-35.5 Mb; **Figure 8**). These loci harbor clusters of *Spin1* and *Spin2,* both members of the spindlin gene family, as well as a large number of genes in the *Btbd35f* family. These regions encompass several gaps on the mm10 mouse reference assembly and are present at variable copy number across the 8 CC founder strains. However, the read depth profile of CC032 at these loci exceeds what is observed in any of the 8 founder strains (**Figure 8; Figure S12**). Several other CC lines exhibit similar read depth patterns across these two X-linked loci, but intriguingly, the strain haplotype origin of these amplified regions is variable (**Figure 8**). CC032 harbors C57BL/6J ancestry across both regions, but strains with 129S1/SvImJ, NOD/ShiLtJ, and PWK/PhJ-derived haplotypes exhibit near identical read-depth signatures (**Figure 8; Figure S12**). This observation would seem to rule out a single common founder effect and imply an incredible rate of structural instability at these loci. However, due to the complex, ampliconic architecture of these loci, we cannot definitively rule out the possibility that assembly, mapping, or genotyping errors have led to misassignment of strain haplotypes in these regions. Long-read sequence data for CC strains may help resolve the architectural complexity of this locus and close standing assembly gaps.

**Figure 8.**
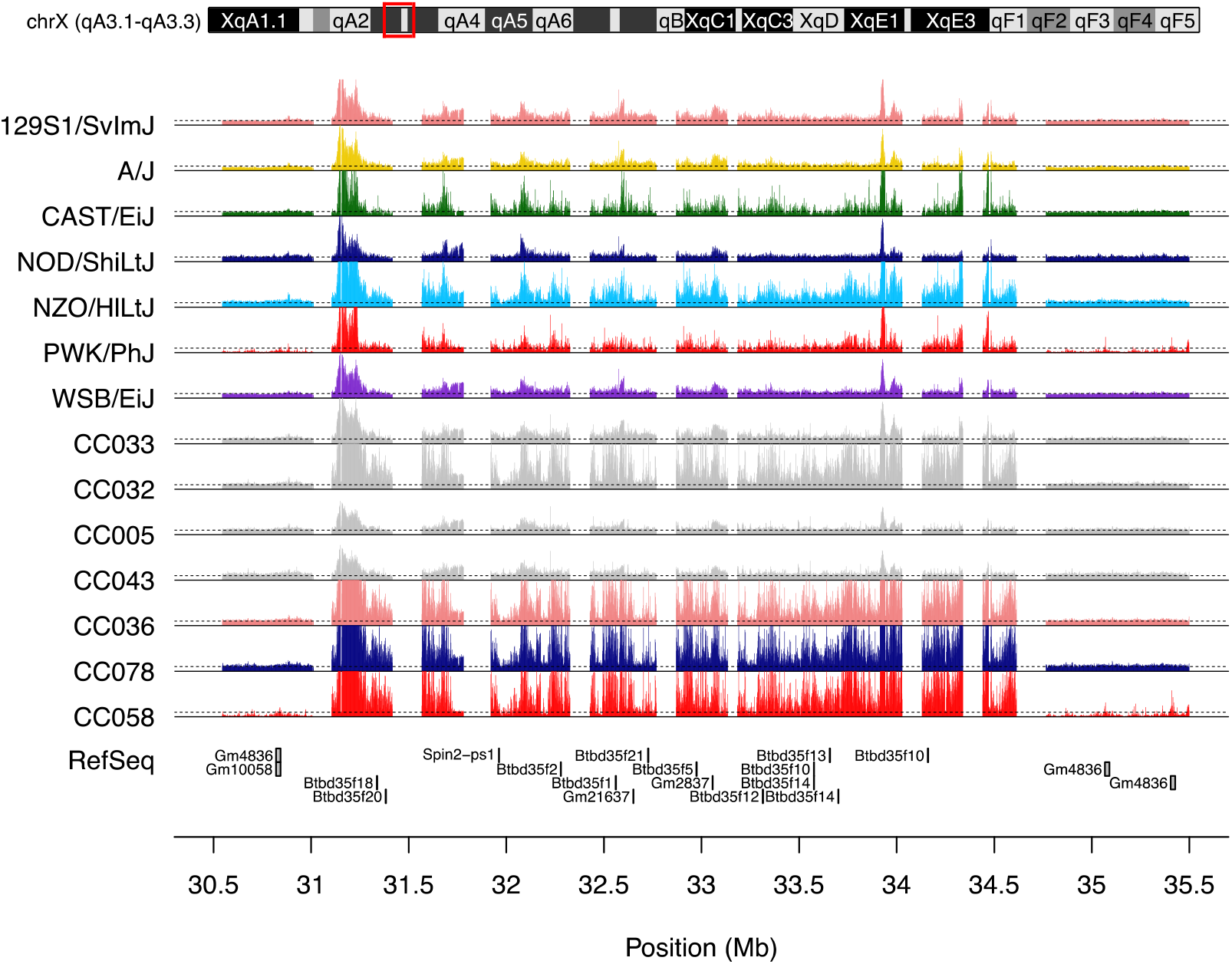
Estimated copy number state in the CC founder strains and 7 CC strains at chrX:30.5-35.5 Mb. Dotted black lines correspond to CN=1. Tracks are color-coded by strain ancestry (A/J: yellow, C57BL/6J: gray, 129S1/SvImJ: pink, NOD/ShiLtJ: dark blue, NZO/HlLtJ: light blue, CAST/EiJ: green, PWK/PhJ: red, and WSB/EiJ: purple). Gaps in the read depth tracks correspond to gaps in the mm10 reference genome assembly. CN states exceeding 10 are clipped for visualization purposes.

Spindlins are chromatin readers that have been shown to directly bind to SLX and SLY (Comptour *et al.* 2014; Kruger *et al.* 2019), although the consequences of this molecular association are poorly understood. Very little is known about *Btbd35f* genes, but they are regulated by SLX/SLXL1- and SLY in spermatids (Moretti *et al.* 2020). Our findings raise the intriguing possibility that the extreme SRD in CC032 is mediated by a complex interplay involving multiple sex-linked ampliconic gene families. However, further work is needed to unravel the potential contributions of spindlins and *Btbd35f* to the dynamic chromatin remodeling during post-meiotic spermatogenesis, and uncover how relative copy number changes at these sex-linked ampliconic genes interface with mechanisms of *Slx*/*Sly*-mediated SRD.

### Sex Ratio Distortion in the Diversity Outbred Mouse Population

The Diversity Outbred (DO) population is a heterogeneous stock developed by outbreeding early generation CC mice from distinct inbreeding funnels (Svenson *et al.* 2012). The population is maintained as 175 outbred families defined by matrilineal inheritance. At every generation, a female from family *A* is mated to a male from a randomly selected family *B*. Their progeny comprise the next generation of DO lineage *A*. As a consequence of this breeding structure, DO mice from a given family are more closely related than DO animals from different families. Thus, if SRD has a genetic basis, males and females from individual DO lineages may sire an excess of males or females relative to Mendelian expectations.

We used published DO breeding records to test for significant SRD in males and females from each of the 175 DO families (**Table S10**; Chesler *et al.* 2016). Females from 15 DO families have sex-biased litters (Binomial Test, uncorrected *P* < 0.05), with 10 families trending toward an excess of males at wean (**Figure 9a**). Similarly, males from 12 DO lineages sire sex-biased litters, with all but three producing an excess of males (**Figure 9b**). For both DO males and females, the observed number of sex-biased families exceeds the ~9 families expected by chance. Sample sizes within each DO lineage are modest (n=67-286; mean = 193), and we are underpowered to detect slight departures from the expected sex ratio. Nonetheless, our findings suggest that the 8-way genotypes associated with DO and CC mice are frequently associated with SRD in both heterozygous and inbred states.

**Figure 9.**
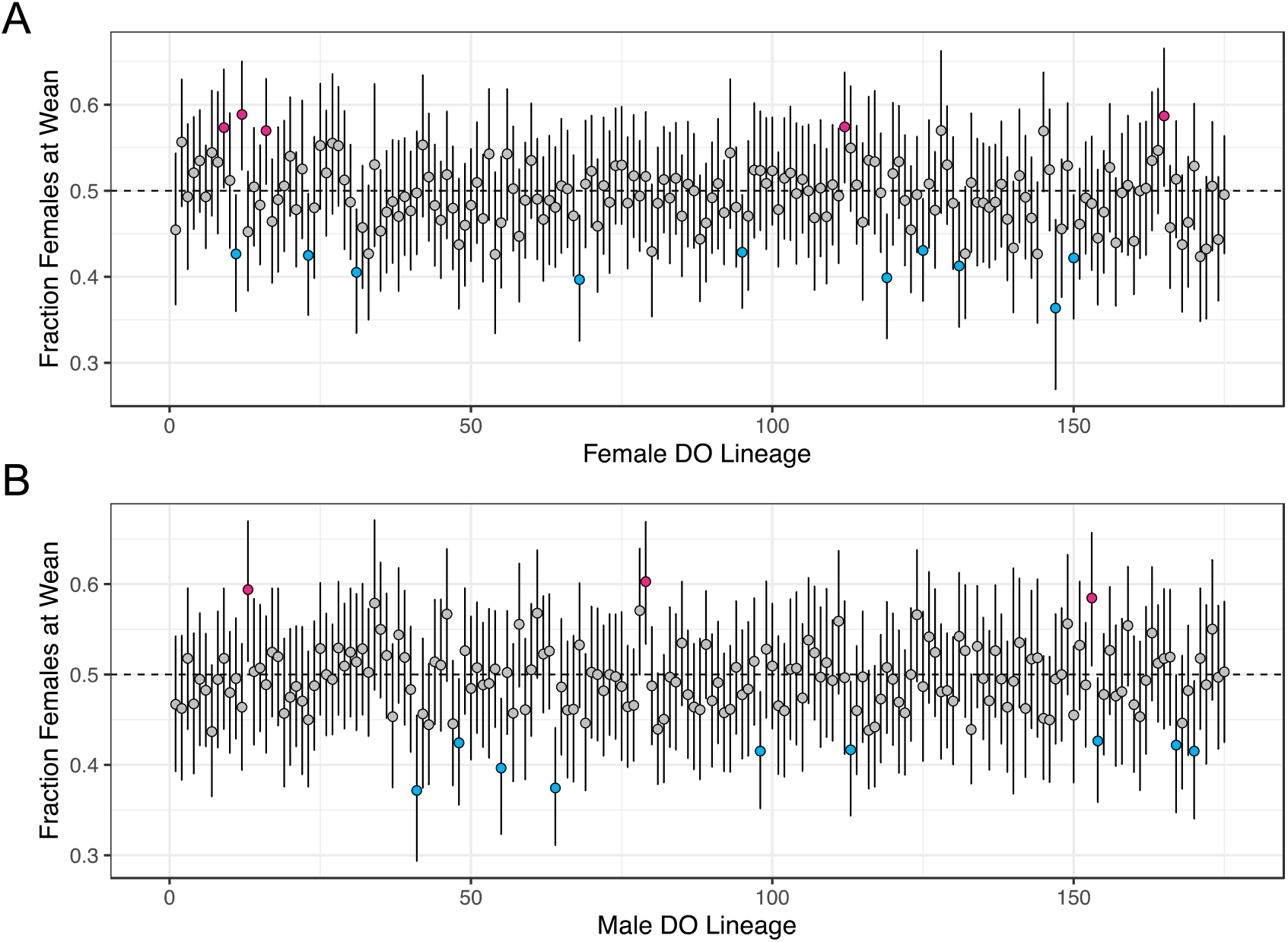
Sex ratios of weaned litters sired by (A) females and (B) males from each DO breeding lineage. Lineages producing significantly male- and female-biased litters are color-coded blue and red, respectively. Error bars correspond to 95% confidence intervals calculated from the binomial distribution.

Despite an excess of both female and male DO lineages with significant SRD, there is no correlation between the sex ratios of progeny sired by dams and sires from a given lineage (Spearman’s Rho = −0.12; *P* = 0.1094; **Figure S13**). Indeed, there are no cases where DO males and DO females from the same lineage both have sex-biased litters (**Figure S13**). Evidently, the biological mechanisms of SRD in the DO (and, potentially, the CC) are largely sex-limited in their manifestation. For instance, germline genetic conflict mediated by sex-linked selfish elements only manifest effects on sex ratios via males whereas maternal effects are only rendered through females.

## DISCUSSION

We have performed the first in-depth analysis of sex ratio distortion in the Collaborative Cross multiparent mouse mapping population. Our integrated analyses of colony breeding records, genome sequences, and phenotypic data uncover widespread SRD in the CC, with several strains exhibiting extreme departures from sex ratio parity. These findings expose the complex basis of sex ratio control, carry important implications for CC husbandry practices, and nominate the CC as powerful resource for studying genetic conflict in action.

We show that CC strain sex ratios are stable over time, consistent across different breeding facilities, and are not broadly dependent on maternal genotype or condition. However, we acknowledge that our analyses are underpowered to detect weak temporal or maternal effects on SRD. We are continuing to compile and analyze breeding records from the CC colony maintained at The Jackson Laboratory, and the addition of new data over time will increase statistical power and may allow detection of smaller effects. It is also critical to emphasize that our investigations are limited to mice reared under a common set of standard laboratory conditions. It remains possible that environmental perturbations or external stressors could induce CC population-wide or strain-specific skews from the sex ratios reported here. Future work is needed to explore the interaction of environmental variables with SRD in the CC. Nonetheless, under the common housing conditions analyzed here, it appears that genetic effects dominant other potential contributors to SRD in the CC population.

Multiple genetic mechanisms can give rise to SRD, including alleles that confer sex differences in embryonic or neonatal survival or genetic mutations that induce sex reversal. We find no consistent relationship between survival in early development and SRD in the CC. Similarly, our analyses of structural variation at key genes in the sex determination pathway rule out sex reversal as a likely explanation for the pervasive SRD in this mapping population. We also show via QTL mapping that there are no large effect single-locus modifiers of SRD segregating in the CC. These results, combined with the general absence of SRD in the inbred parental CC lines and their F1 hybrids (Shorter *et al.* 2019a), imply that the underlying genetic mechanisms of SRD are most likely attributable to multilocus combinations of alleles that are only revealed in the 8-way CC population.

These considerations lead us to speculate that the SRD in this population arises, at least in part, from the de-coupling of cryptic selfish sex-linked drive elements and their co-evolved suppressors. Several lines of evidence lend support to this hypothesis. First, such genetic conflicts necessary involve the interaction of multiple loci, aligning with the absence of SRD in the CC founder strains and the lack of any large, single-locus effects in our QTL mapping. Second, we observe minimal evidence for maternal effects and maternal genotype-dependence on SRD, implying that SRD in the CC is most frequently determined by mechanisms rendered through the male germline. Conflict between feuding drive elements on the X and Y chromosomes is, by genetic necessity, limited in expression to males. Third, transmission distorters typically act by disabling or killing non-carrier gametes, imposing a fitness cost to carriers (Zanders and Unckless 2019). Many sex-biased CC strains are characterized by small litter sizes (**Figure 3a**) and markers of reduced male fertility, including low testis weights, reduced sperm density, and impaired sperm motility (**Figure 7**).

One compelling candidate system is the SYCP3-like family of ampliconic sex-linked transmission distorters: *Slx* and *Slxl1* on the X chromosome and their Y-linked paralog, *Sly*. Prior work has demonstrated that genetic imbalance of SLX/SLXL1 and SLY leads to disruption of the gene regulatory program in spermatids, sex ratio distortion, and infertility in house mice (Cocquet *et al.* 2009, 2012; Kruger *et al.* 2019). The 8 CC founder strains differ in their native copy number status at these genes (Morgan and Pardo-Manuel de Villena 2017), and many sex-biased CC lines have inherited a relative excess or deficit of *Slx/Slxl1* gene copies relative to *Sly*, consistent with their observed SRD.

However, we do not find a simple, overall relationship between SRD and *Slx/Slxl1:Sly* CN ratio across this population and several strains with extreme *Slx/Slxl1:Sly* ratios do not exhibit expected patterns of SRD (**Figure 5A**). Some genomic copies of these genes may be nonfunctional, and future work to quantify their mRNA or protein abundance in CC strains may help clarify the presumed underlying relationship between the protein products of these ampliconic genes and SRD. Beyond this possibility, recent work has begun to uncover the complexity of *Slx/Slxl1* and *Sly* mediated conflict, suggesting that a simple linear relationship between gene copy number ratio and the magnitude of SRD is potentially overly simplistic. In particular, SLX/SLXL1 and SLY compete for binding to members of a second sex-linked amplicon gene family – the spindlin proteins SSTY1/2 and SPIN1 (Kruger *et al.* 2019; Moretti *et al.* 2020) – to antagonistically regulate large numbers of sex-linked and autosomal genes in post-meiotic spermatids. It is noteworthy that many sex-biased CC strains also have extreme spindlin genomic copy numbers (**Figure 5C**). Additionally, our focused investigations in CC032/GeniUncJ reveal apparent *de novo* expansions at two spindlin gene clusters on chrX, seemingly implicating these genes in the extreme SRD that characterizes this strain. We speculate that spindlins, and potentially other sex-linked ampliconic genes, interact with *Slx/Slxl1* and *Sly* to modulate the strength of SRD in different CC strains. Future work to probe patterns of differential gene regulation in round spermatids from strains with extreme SRD versus those siring sex-balanced litters may help unlock the complex molecular mechanisms of SRD in this population.

Overall, the widespread trend of SRD across the CC and DO mouse populations seems to suggest that the intersubspecific 8-way genotypes segregating in these populations unmask a complex network of segregation distorters that are silenced in the context of individual inbred strains. Here, we focused on sex ratio distortion, as phenotypic sex provides a faithful readout of chromosome transmission (assuming no sex reversal). However, the high frequency of SRD across these populations raises the parallel prospect that selfish elements on other chromosomes could bias transmission in these 8-way diverse mouse populations. Indeed, a segregation distorter on chr2qC3 was previously identified via routine genetic monitoring in the DO population (Didion *et al.* 2015; Chesler *et al.* 2016). Specifically, the WSB/EiJ allele at this locus exhibits preferential segregation to the maternal oocyte during asymmetric female meiosis and threatened to drive to fixation, purging segregating variation at this locus from the DO. Although the DO maintenance breeding program is designed to minimize the potential for drive (Chesler *et al.* 2016), it is expected that, overtime, *de novo* evolution or the recombination of existing drive elements onto permissible genetic backgrounds could allow transmission distorters to take root.

Our findings also carry practical implications for CC strain maintenance and experimental design. Many CC strains exhibit only slight or no departure from Mendelian sex ratio expectations, but several yield strongly sex-biased litters. CC065/UncJ and CC032/GeniUncJ are the most notable examples, with just one male and one female in every three live-weaned pups, respectively. Given the modest reproductive output of most CC lines, these aspects of strain reproductive performance highlight the need for implementing strain-specific breeding programs to maintain stable colonies. Such practices are already in place at The Jackson Laboratory to ensure the long-term shelf-stability of this important diverse mouse resource, but should also be embraced in the settings of individual laboratories. Importantly, these breeding challenges are not necessarily eliminated by outcrossing, as we document several CC-RIX backgrounds with SRD (**Figure S6**) and observe an excess of sex-biased lineages in the Diversity Outbred mapping population (**Figure 9**).

The CC population is an established resource for complex trait mapping and systems genetics investigation, and has yielded powerful new mouse models of human disease (Churchill *et al.* 2004; Aylor *et al.* 2011; Philip *et al.* 2011; Rogala *et al.* 2014; Srivastava *et al.* 2017; Green *et al.* 2017). At the same time, the CC panel represents a pedigreed, well-resourced population optimally suited for investigations into the fundamental properties of genetic inheritance, including chromosome transmission. Each realized CC line harbors a unique multilocus combination of haplotypes from three cardinal house mouse subspecies, providing a real-time window into how intersubspecific allele permutations shape genome function and evolution. Our work has spotlighted the CC as a uniquely powerful platform for studying intragenomic genetic conflict and SRD in house mice and lays the groundwork for future investigations into the molecular basis of this important biological phenomenon.

## ACKNOWLEDGEMENTS

We gratefully acknowledge the contribution of the Histopathology and Microscopy Scientific Services at The Jackson Laboratory for expert assistance with the work described in this publication. We are also indebted to Racheal Wallace for conserving cage cards from all Collaborative Cross mating units, enabling us to compile comprehensive breeding records from this mouse population. We thank Candice Baker and Cat Lutz for sharing breeding data from CC-RIX crosses. This work was funded by a NIGMS MIRA (R35GM133415) awarded to BLD. FB was supported by a Research Experience for Undergraduate Site Award (DBI-1262049; PI: Bob Braun).

---

**Figure S1.**
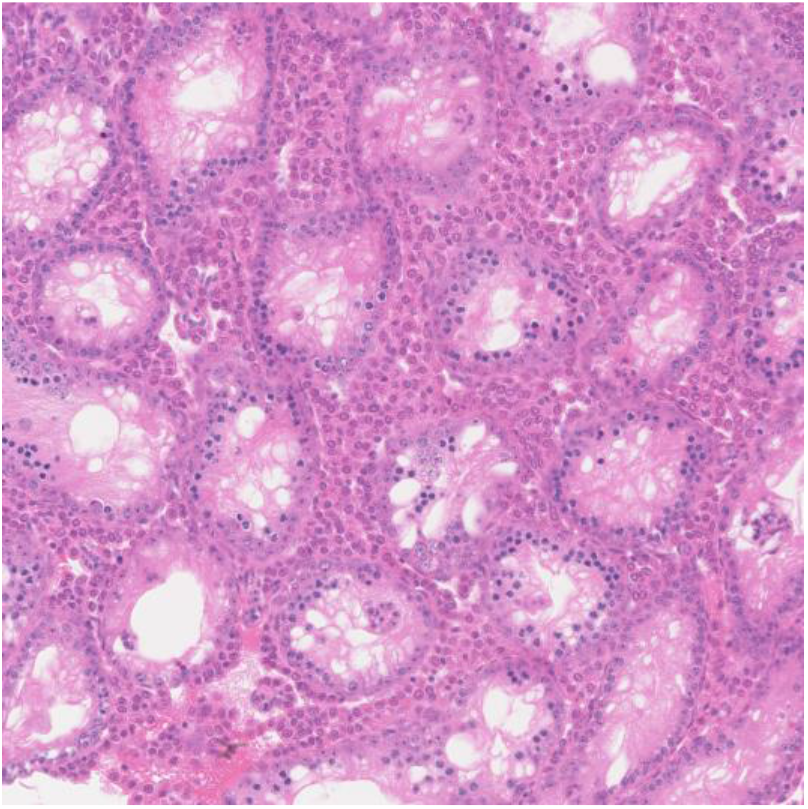
Hematoxylin and Eosin-Y stained testis cross section from a CC032/GeniUncJ male at ~40x magnification. A high proportion of seminiferous tubules exhibit large vacuoles and germ cell loss.

**Figure S2.**
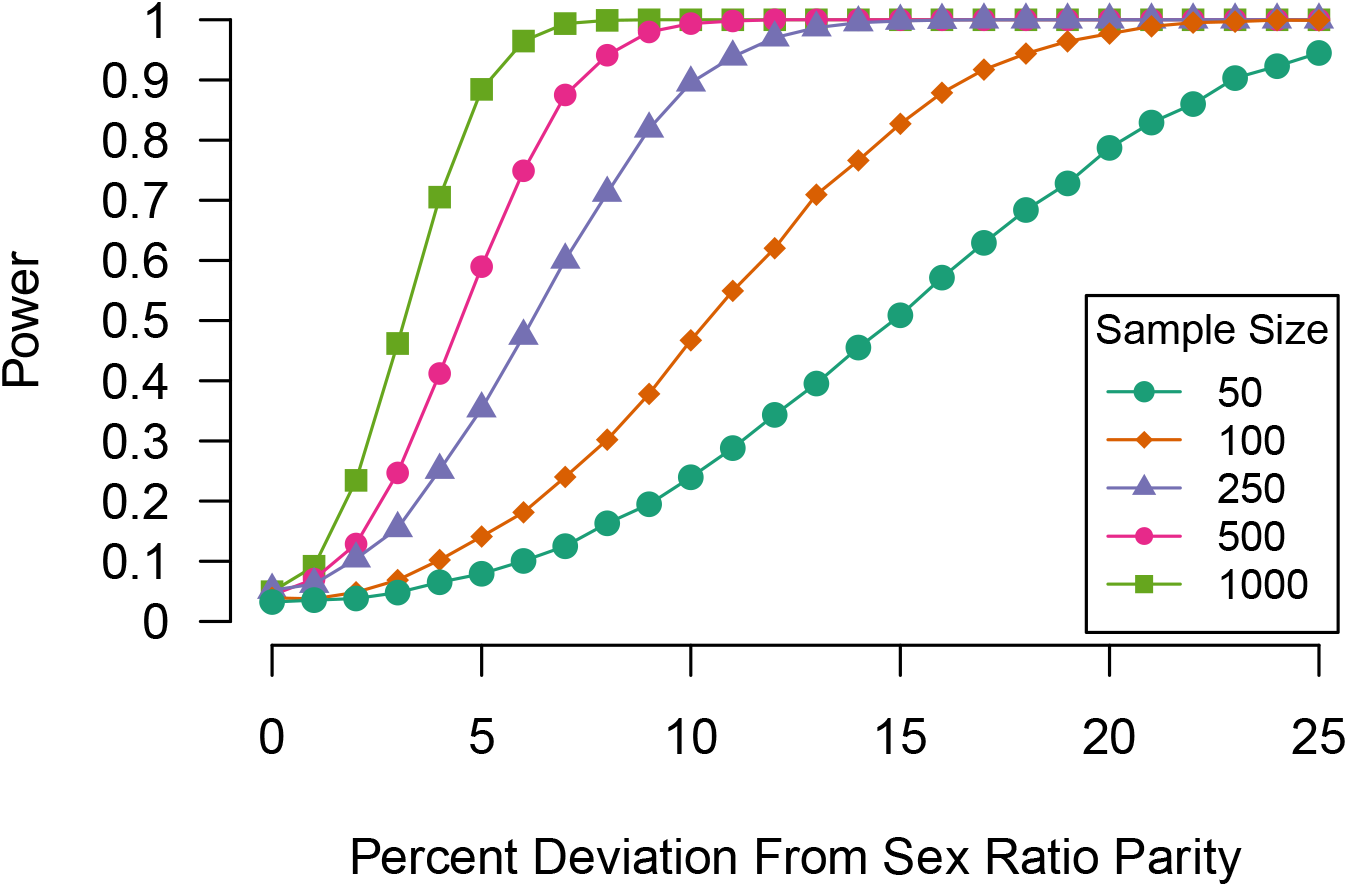
Statistical power to detect sex ratio distortion using a two-way binomial test. Colors and plotting shapes denote different sample sizes.

**Figure S3.**
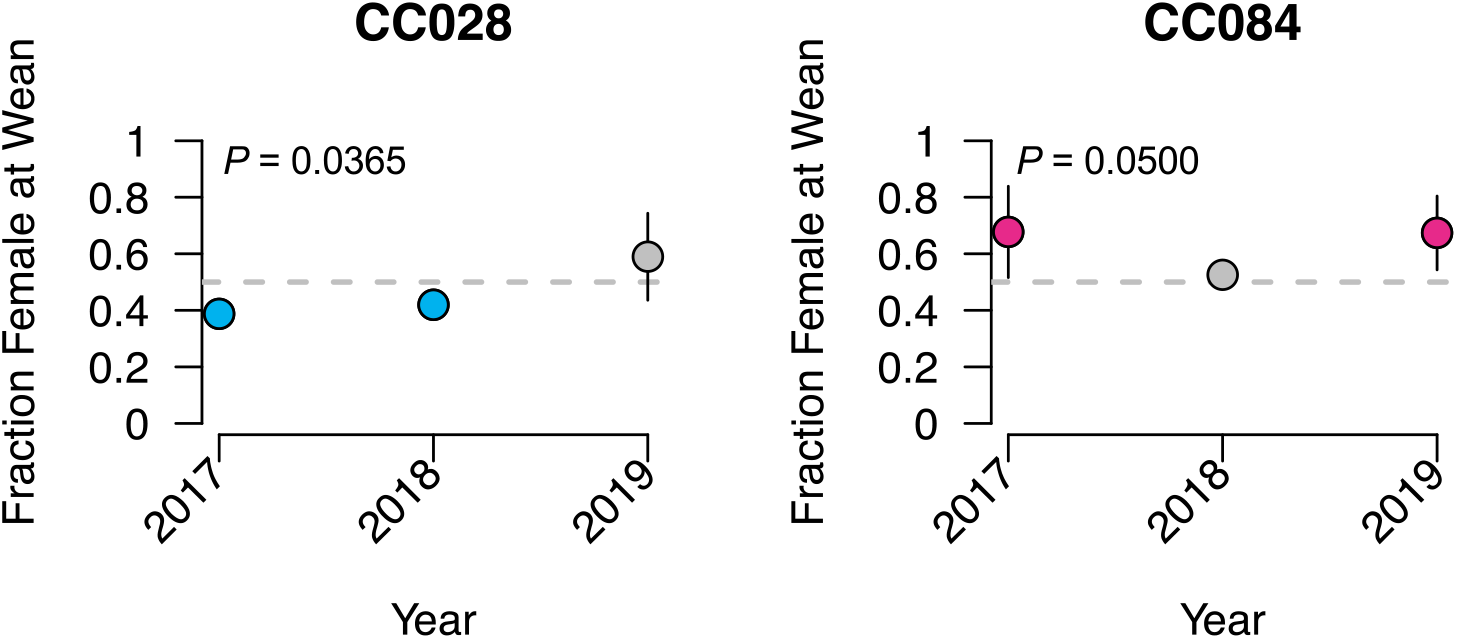
The fraction of females at wean exhibits mild variation over time for CC028/GeniUncJ and CC084/TauJ. Error bars correspond to 95% binomial confidence intervals. Sex ratios are estimated with high precision for some strain-year combinations, and confidence intervals are masked by the plotting characters. Samples with significant male- and female-sex biases are color coded blue and red, respectively.

**Figure S4.**
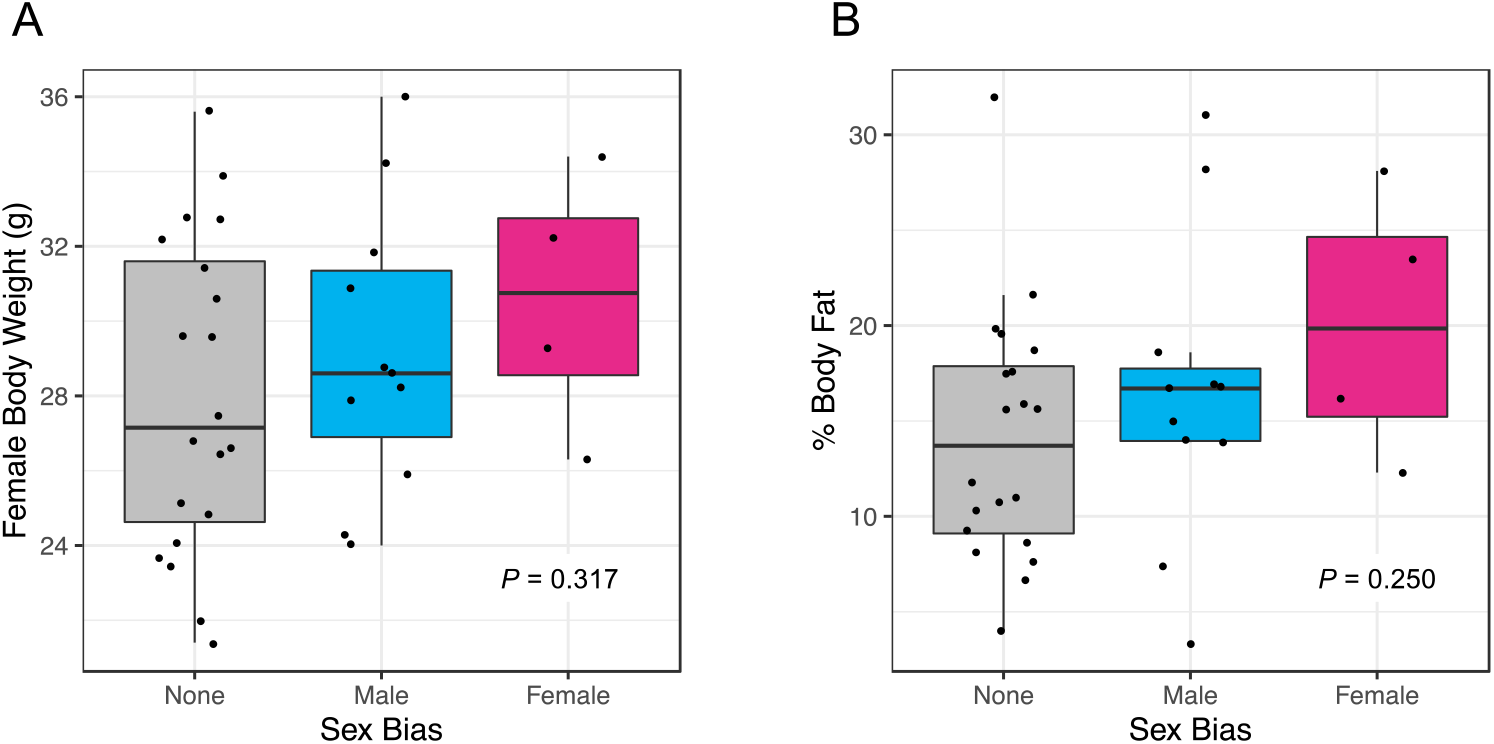
Relationship between sex ratio bias and mean adult female (A) body weight and (B) body fat percentage. Each point corresponds to a single CC strain. CC strains are designated as male- or female-biased based according to per-strain binomial tests *(P* < 0.05).

**Figure S5.**
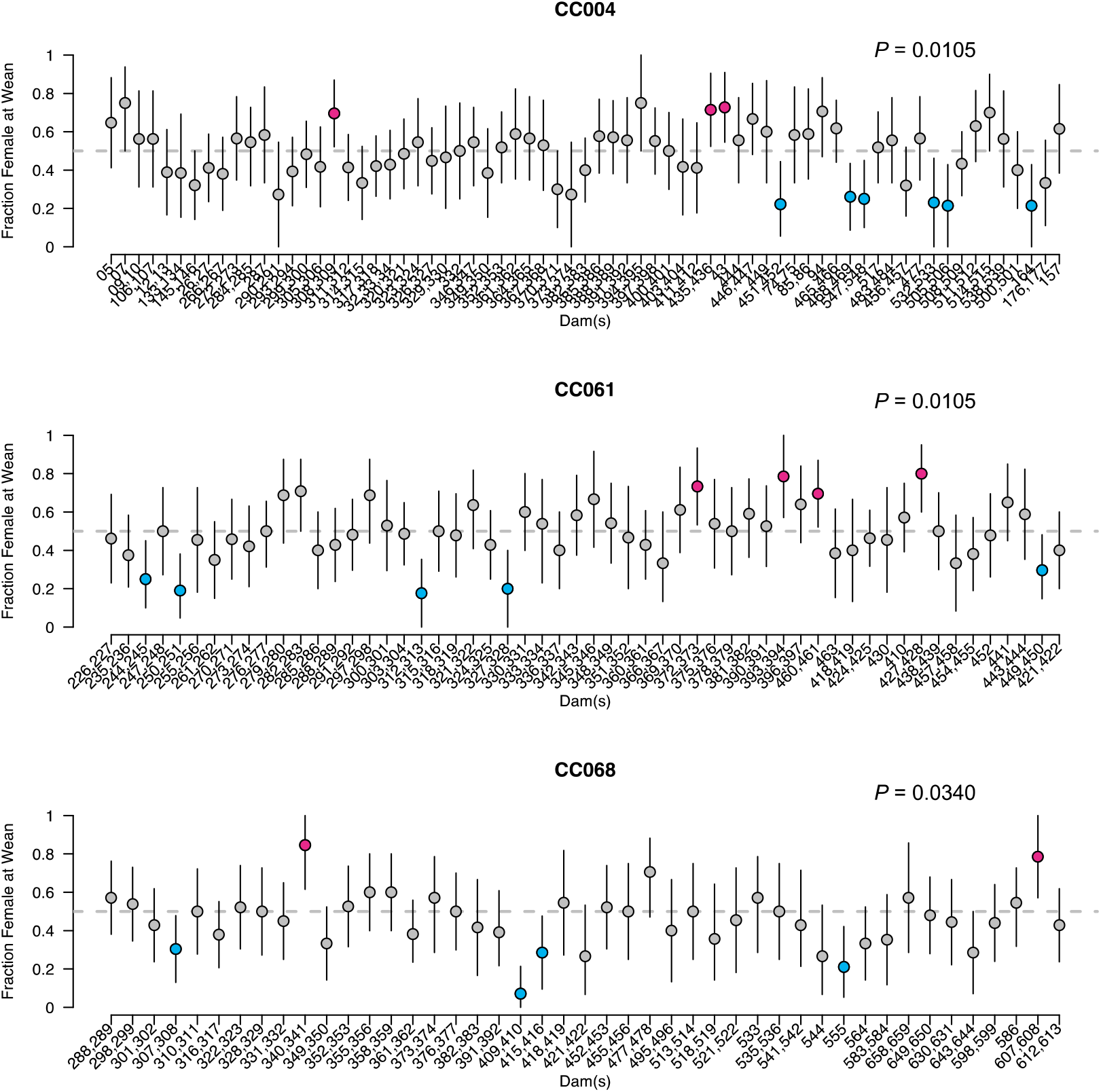
Dam identity exerts a weak influence on offspring sex ratios in CC004/TauUncJ, CC061/GeniUncJ, and CC068/TauUncJ. For each strain, the sex ratio of animals sired by each dam (or pair of dams, in the case of trio matings) is plotted as the fraction of females at wean. Dams producing significantly female- or male-biased litters are denoted by the red and blue points, respectively. Error bars correspond to 95% confidence intervals calculated from the binomial distribution.

**Figure S6.**
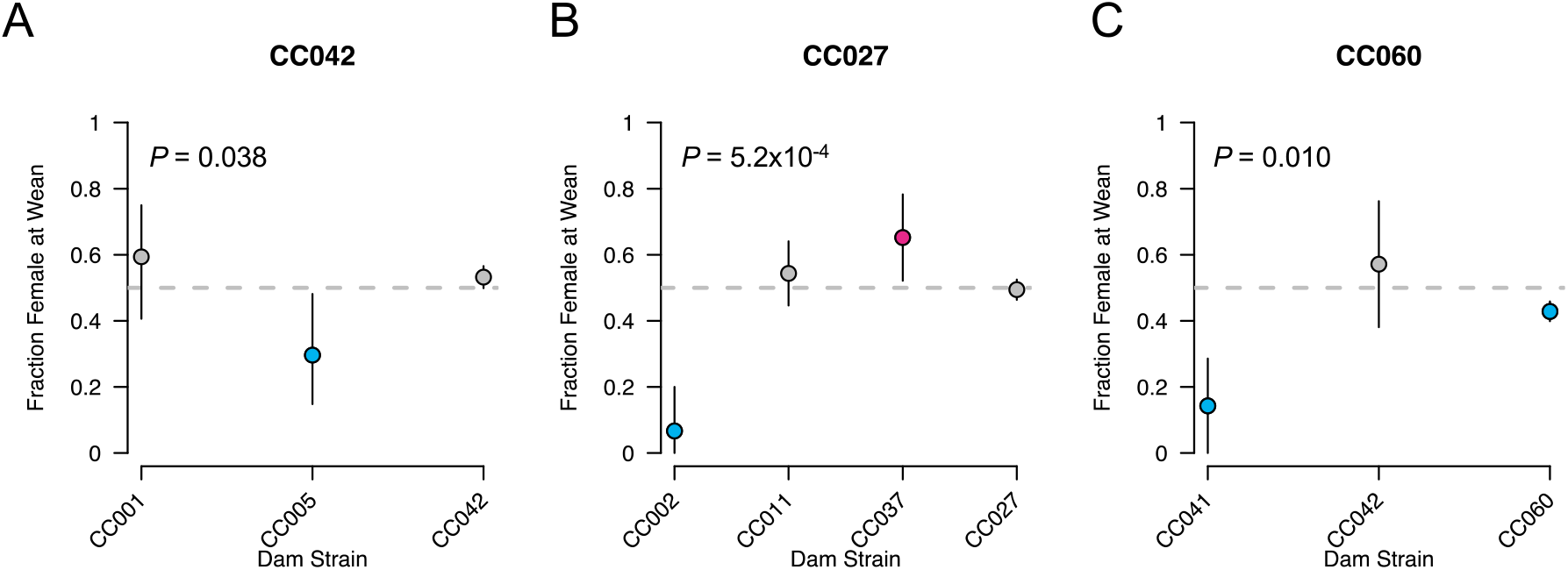
CC males from strains (A) CC042/GeniUncJ, (B) CC027/GeniUncJ, and (C) CC060/UncJ sire litters with variable sex ratios depending on the genetic background of the dam. Error bars correspond to 95% confidence intervals calculated from the binomial distribution. Dam strains producing significantly male- or female-biased litters are color-coded blue and red, respectively.

**Figure S7.**
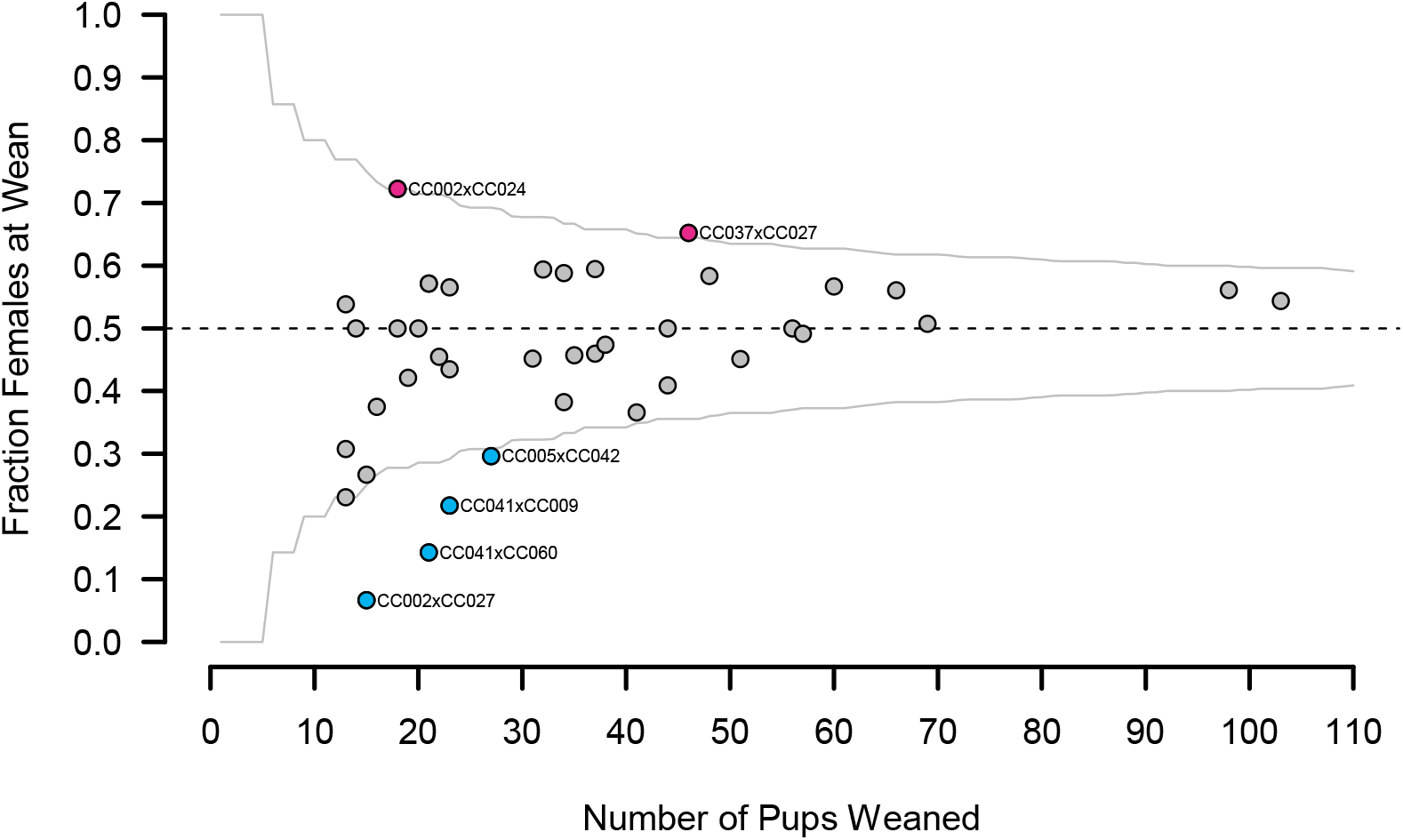
Sex ratio distortion in CC-RIX mice. The horizontal dashed line corresponds to the expected 1:1 sex ratio. The gray solid curves delimit the range of sex ratio variation expected due to binomial sampling for a given sample size. Crosses that yield significantly female- or male-biased litters are shown in red and blue, respectively. Point labels specify the associated CC-RIX cross in the format: dam x sire.

**Figure S8.**
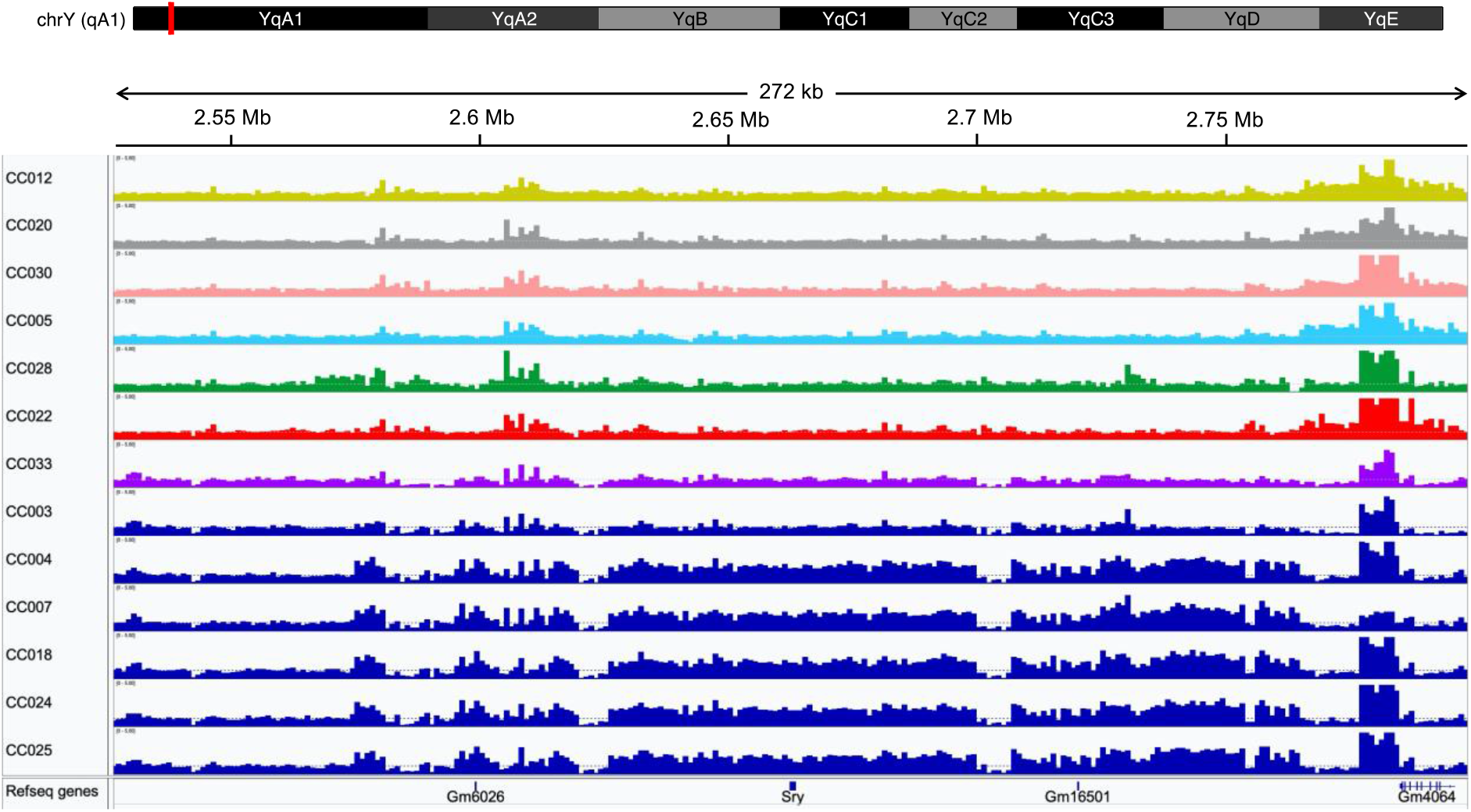
Estimated genomic copy number in 1kb windows across the *Sry* locus on the short arm of chrY for a sample of CC lines. CC lines carrying the NOD/ShiLtJ Y chromosome are depicted in dark blue. Other CC strains are color coded by the parental origin of their Y chromosome: yellow (A/J), gray (C57BL/6J), pink (129S1/SvImJ), light blue (NZO/HlLtJ), green (CAST/EiJ), red (PWK/PhJ), and purple (WSB/Eij) The faint horizontal dashed line on each plot corresponds to the expected CN=1 state.

**Figure S9.**
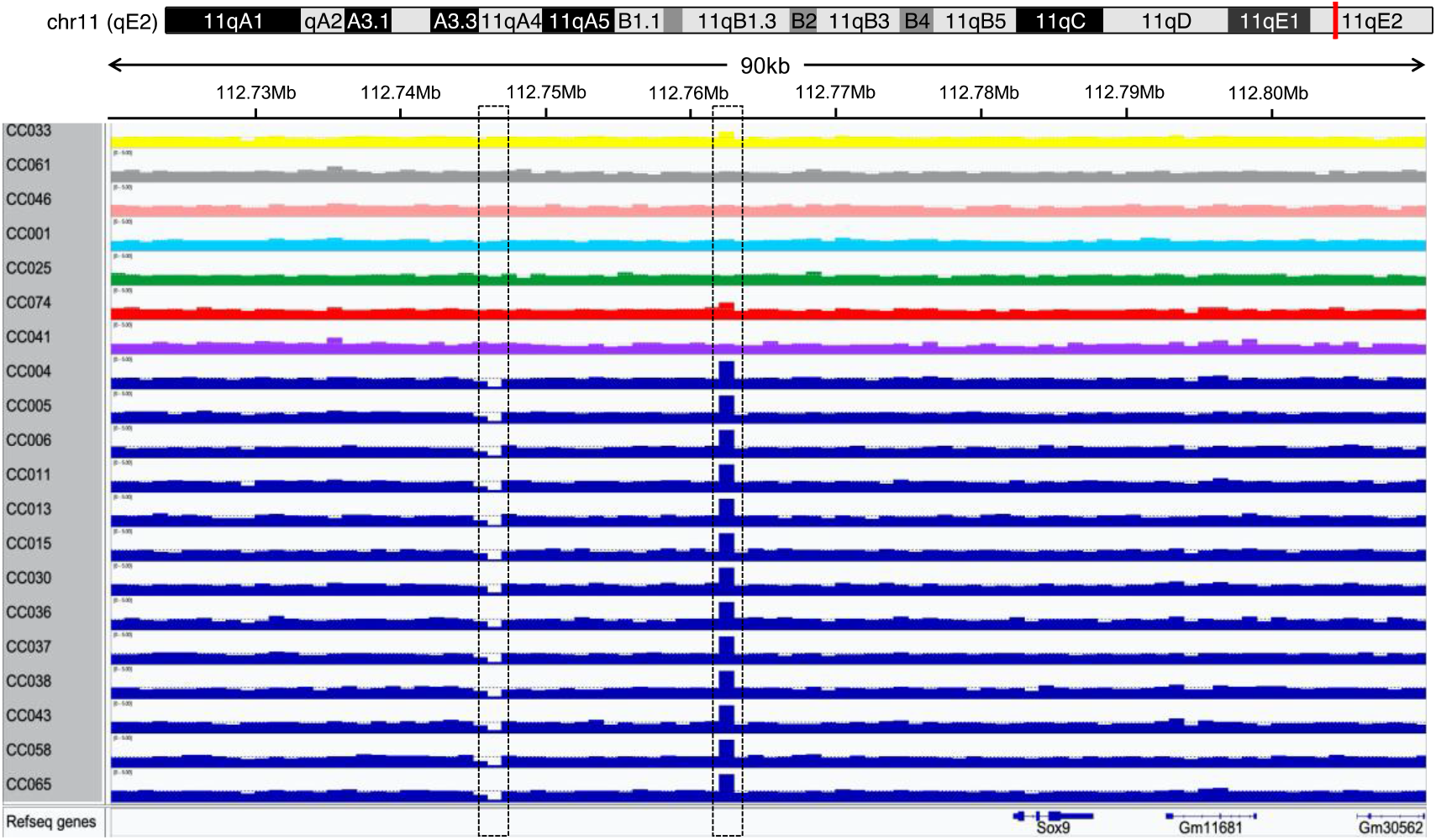
Estimated genomic copy number in 1kb windows across the *Sox9* locus on chr11 for a sample of CC lines. CC lines carrying the NOD/ShiLtJ haplotype at this locus are depicted in dark blue. Other CC strains are color coded by the parental origin of their chromosome: yellow (A/J), gray (C57BL/6J), pink (129S1/SvImJ), light blue (NZO/HlLtJ), green (CAST/EiJ), red (PWK/PhJ), and purple (WSB/EiJ) The horizontal dashed line on each plot corresponds to the expected CN=2 state. Dashed black boxes highlight a small deletion and duplication in the *Sox9* distal upstream regulatory region that are specific to the NOD/ShiLtJ haplotype.

**Figure S10.**
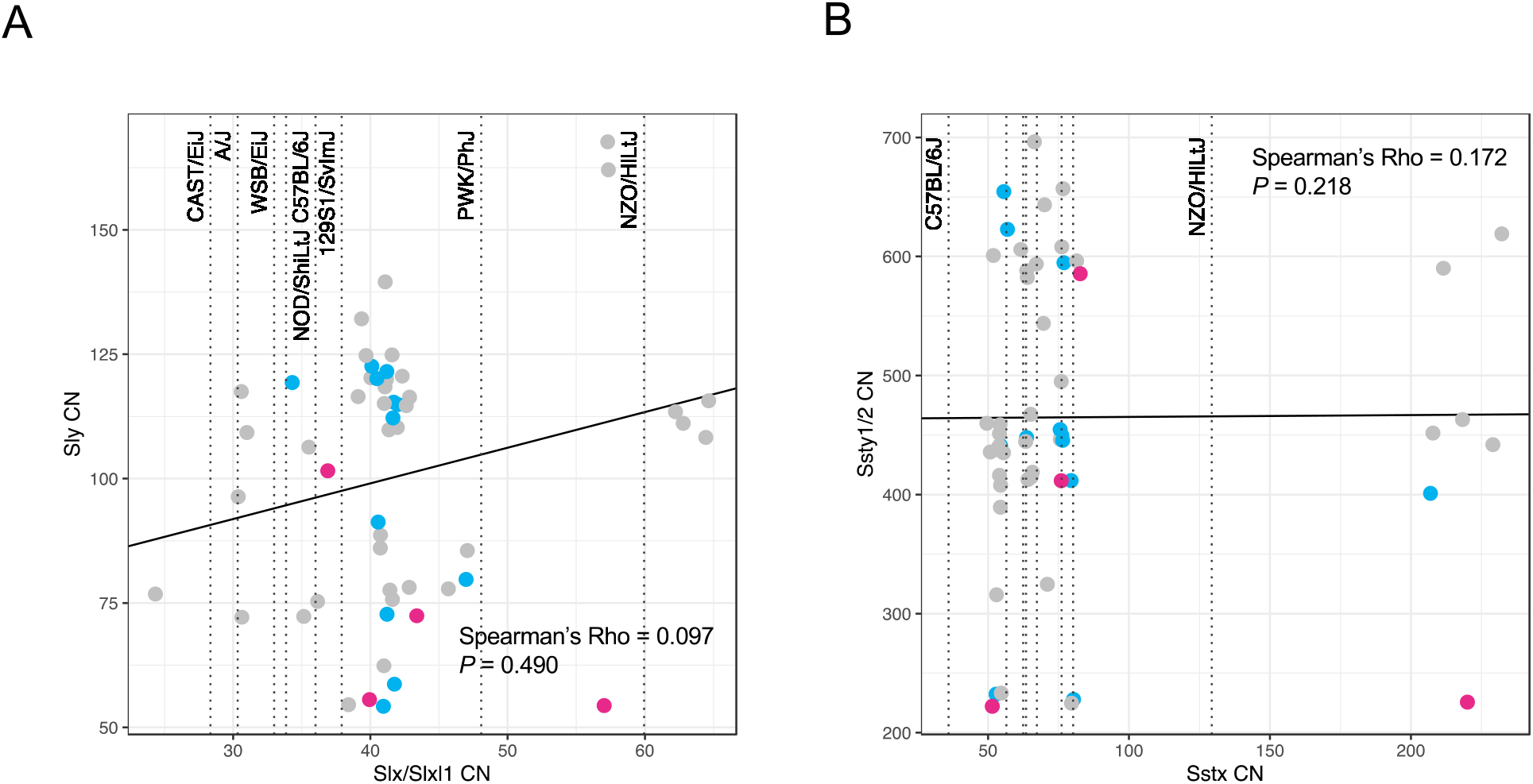
(A) *Slx/Slxl1* and *Sly* copy number estimates for the CC strains. Vertical dashed lines denote estimated *Slx/Slxl1* copy numbers for each of the 8 CC founder strains. The solid black line is the least squares trend line fit to the data. (B) *Sstx* and *Ssty1/2* copy number estimates for the sequenced CC strains. As in (A), vertical dashed lines indicate the estimated *Sstx* copy number state of the inbred CC founder strains and the solid black line is an overall trend line fit to the data using the method of least squares. Strains with significantly male-and female-biased sex ratios are color-coded blue and red, respectively.

**Figure S11.**
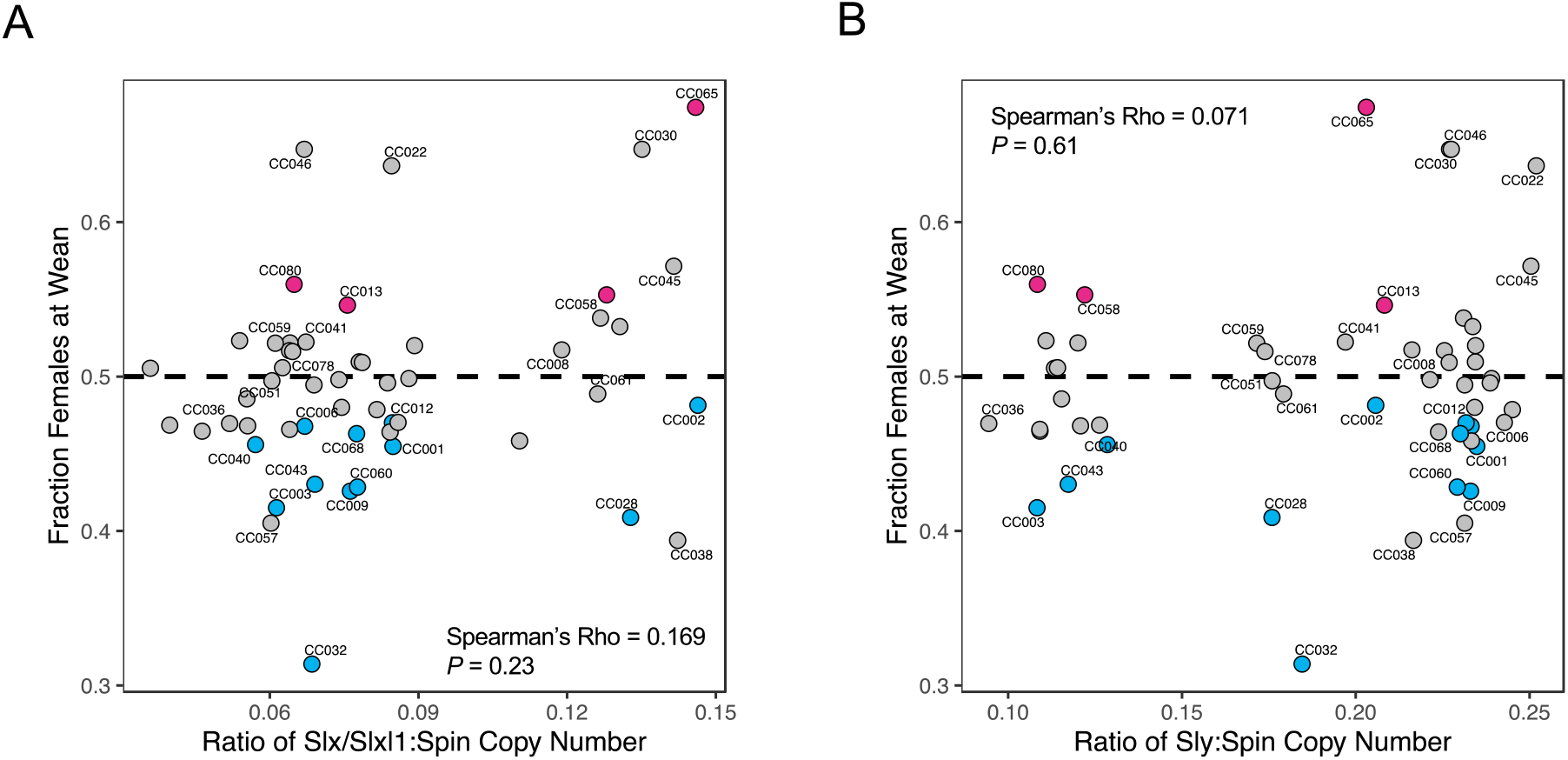
(A) Relationship between the ratio of *Slx/Slxl1* and spindlin copy numbers and SRD and (B) the relationship between *Sly* and spindlin copy number and the fraction of females at wean in the CC panel. Points corresponding to significantly sex-biased strains are color-coded (blue = male-biased; red = female-biased). Horizontal dashed black line corresponds to a balanced sex ratio of 0.5.

**Figure S12.**
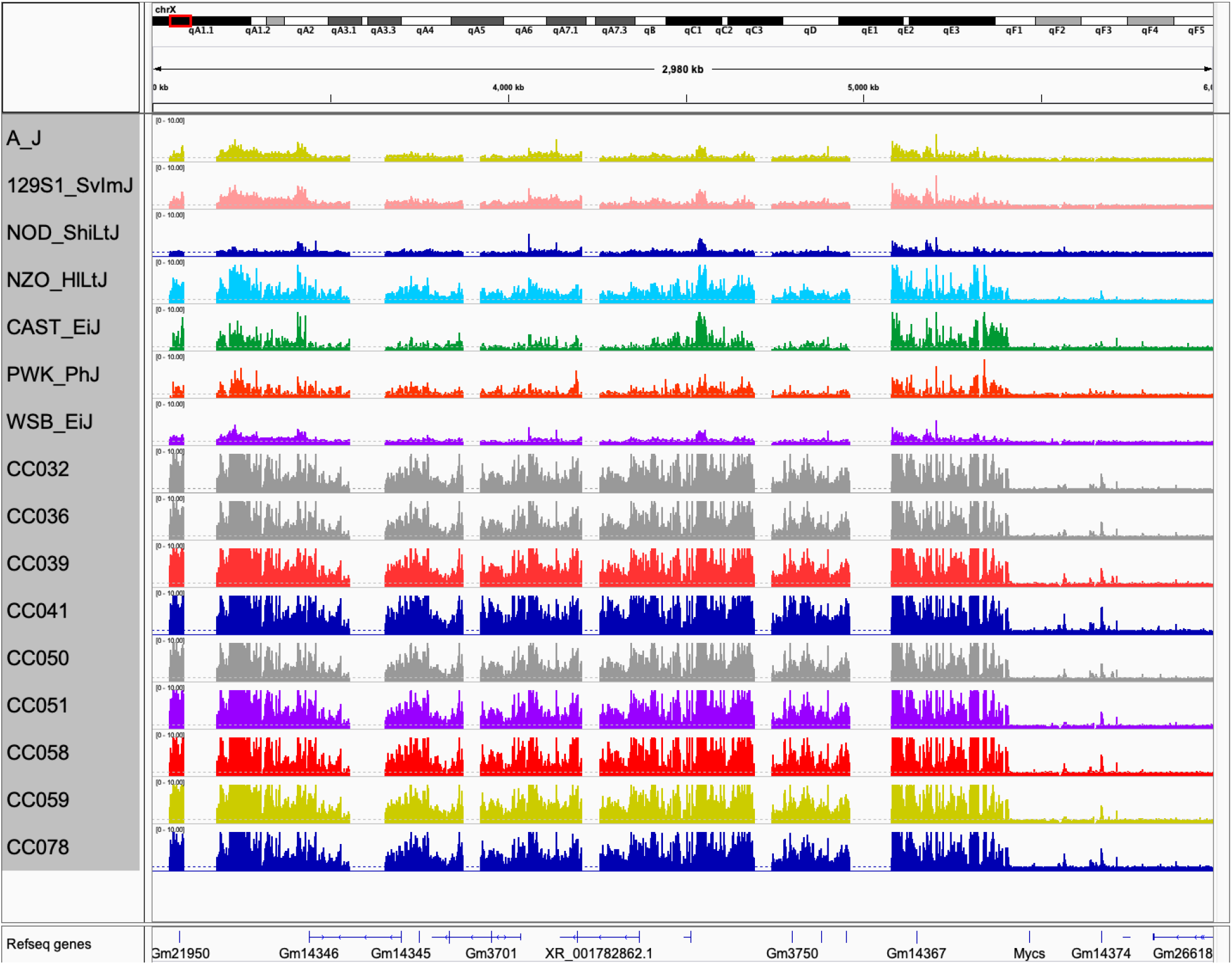
Estimated copy number state in the CC founder strains and 7 CC strains at chrX:3-6Mb. Dotted black lines correspond to CN=1. Tracks are color-coded by strain ancestry (A/J: yellow, C57BL/6J: gray, 129S1/SvImJ: pink, NOD/ShiLtJ: dark blue, NZO/HlLtJ: light blue, CAST/EiJ: green, PWK/PhJ: red, and WSB/EiJ: purple). Gaps in the read depth tracks correspond to gaps in the mm10 reference genome assembly. CN states exceeding 10 are clipped for visualization purposes.

**Figure S13.**
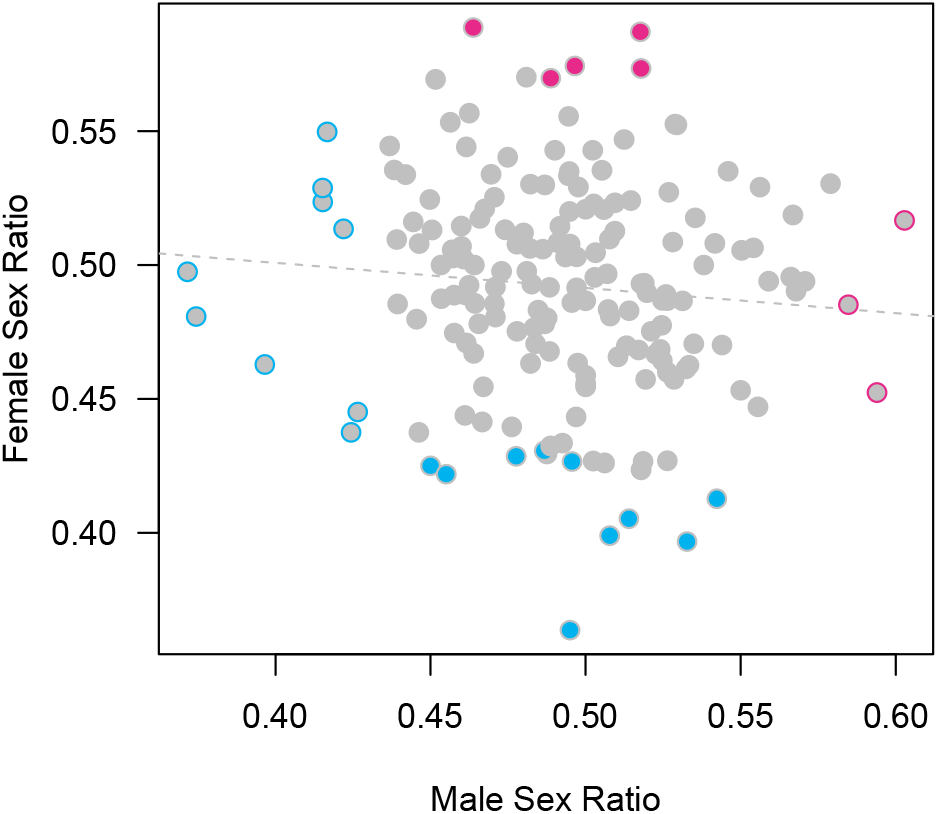
Correlation between the sex ratio of litters sired by males and females from the same DO breeding lineage. The outline color of each point denotes the direction of any significant sex bias associated with male DO lineages. Conversely, the solid color of each point indicates the direction of significant sex bias associated with female lineages. No lineages exhibit both male and female-associated sex bias. Blue: male bias; Red: female bias. Gray: no significant sex bias.

